# Structural proteomics defines a sequential priming mechanism for the progesterone receptor

**DOI:** 10.1101/2024.09.06.611729

**Authors:** Matthew D. Mann, Min Wang, Josephine C. Ferreon, Michael P. Suess, Antrix Jain, Anna Malovannaya, Roberto Vera Alvarez, Bruce D. Pascal, Raj Kumar, Dean P. Edwards, Patrick R. Griffin

## Abstract

The progesterone receptor (PR) is a steroid-responsive nuclear receptor with two isoforms: PR-A and PR-B. Disruption of PR-A:PR-B signaling is associated with breast cancer through interactions with oncogenic co-regulatory proteins (CoRs). However, molecular details of isoform-specific PR-CoR interactions remain poorly understood. Using structural mass spectrometry, we investigate the sequential binding mechanism of purified full-length PR and intact CoRs, steroid receptor coactivator 3 (SRC3) and p300, as complexes on target DNA. Our findings reveal selective CoR NR-box binding by PR and unique interaction surfaces between PR and CoRs during complex assembly, providing a structural basis for CoR sequential binding on PR. Antagonist-bound PR showed persistent CoR interactions, challenging the classical model of nuclear receptor activation and repression. Collectively, we offer a peptide-level perspective on the organization of the PR transcriptional complex and infer the mechanisms behind the interactions of these proteins, both in active and inactive conformations.

## Introduction

The progesterone receptor (PR) is a steroid-activated nuclear receptor and belongs to the subfamily of steroid hormone receptors (SRs) that includes the estrogen receptor, androgen receptor, mineralocorticoid receptor, and glucocorticoid receptor. The SRs, including PR, exhibit remarkable functional diversity in mediating cell/tissue and target gene-specific responses, largely driven by conformational dynamics of the protein that enables its binding to DNA response elements and unique subsets of co-regulatory proteins (CoRs). Like other SRs, PR is a modular protein composed of well-folded ligand binding (LBD) and DNA binding (DBD) domains that are connected via a structurally dynamic hinge region, and an intrinsically disordered (ID) N-terminal domain (NTD). The hinge, additionally termed carboxyl terminal extension (CTE), is more than a flexible linker. It forms an extended loop that interacts with the minor groove of DNA flanking either side of inverted repeat progesterone response element (PREs) to extend the protein-DNA interface beyond that of the core DBD that binds major groove of HREs.^1, 2^ SRs also contain two transcriptional activation functions (AFs), ligand-independent AF-1 in the NTD and ligand-dependent AF-2 in the LBD, that provide interaction surfaces for CoRs.^3–5^

PR is important for development, proliferation and differentiation of female reproductive tissues during the reproductive cycle and pregnancy.^6–11^ The biological response to progesterone is mediated by two distinct protein isoforms: PR-A and PR-B, that are expressed through alternate utilization of two promoters of the same gene. PR-A is an N-terminal truncation (missing aa 1-164) of full-length PR-B, with each having distinct cell/tissue-dependent physiological roles. Generally, PR-A is a weaker transcriptional activator than PR-B, and both isoforms are typically co-expressed in equal proportions in most normal tissues. However, their ratios have been reported to be highly variable in pathological conditions. Both PR isoforms contribute to the onset and tumorigenesis of breast cancer, and increased expression of PR-A has been shown to alter progestin responsiveness in cancer phenotypes.^8, 12–15^ Interestingly, known CoRs are shared between isoforms, yet each isoform’s transcriptional activity differs. The mechanistic basis for differences in the activity of the isoforms remains unclear but is generally believed to be due to differences yet to be determined in structural conformations and, thereby, CoR interactions.^10, 16–18^

To better understand PR biology requires detailed structural analysis of the full-length PR isoforms, associated with full-length CoRs: steroid receptor coactivator 3 (SRC3) and histone acetyltransferase p300 as a complex on target DNA, and an understanding of how protein interactions within the complex and structural conformations can affect the activity of PR. The conformational flexibility of SRs and CoRs, coupled with their large sizes, make them unsuitable for either high-resolution NMR or X-ray crystallography analysis. Recent low-resolution cryoEM studies using PR:SRC2:p300 complex also failed to provide details of specific interactions of PR with CoRs at the atomic level, thus limiting the utility of these analyses.^19^ Here, we present an amino-acid resolution view of canonical PR-A and PR-B transcriptional complex, including SRC3 and p300, on PR target DNA. High resolution crystal structures have been determined for the PR DBD-CTE^1^ complexed with a consensus inverted repeat palindromic progesterone response element (PRE) DNA, and the PR LBD complexed with a progesterone agonist^20^ or antagonists with a peptide from a transcriptional co-repressor.^21, 22^ However, there is a lack of structural detail for full-length PR and CoRs as a complex with PRE DNA.^21, 23–25^

An alternative to classical structural techniques is structural proteomics. Hydrogen-deuterium exchange (HDX) works on the principle of backbone amide exchange, where protein backbone amide hydrogens will freely exchange to deuterium upon regular protein motion.^26–30^ This is a useful technique where differential deuterium exchange informs protein conformational changes and protein-protein interaction sites. Similarly, crosslinking (XL) coupled with mass spectrometry (MS) provides information on amino acid proximity, which is useful in determining protein-protein interaction regions.^31–34^ Through a combination of HDX-MS and XL-MS, we present a higher-resolution structural understanding of the organization of the PR ternary complex compared to available structures. These data show that PR interacts with SRC3 and p300 in an isoform-specific and ligand-specific manner that may impart distinct SR function. The structural information gained from our studies is essential for advancing our understanding of the molecular mechanism of action of PR and other SRs.

## Results

### Strep-II tagged recombinant PR and CoRs generates stable protein and complexes

Intact full-length PR and CoRs were expressed and purified as recombinant proteins in Sf9 insect cells using the baculovirus system.^35^ This is an ideal expression system for full-length PR shown previously to retain native folding and post-translational modifications including phosphorylation on the same sites as occurs with endogenous PR in mammalian cell types.^36, 37^ Additionally purified full-length PR from the baculovirus system was demonstrated to exhibit stoichiometric ligand binding activity, high affinity binding to specific progesterone response element (PRE) DNA, including free DNA and when assembled on nucleosomes, and transcriptional activity in cell-free assays.^35, 38–41^ To refine the previous methods for expression and purification of PR, a Strep-II affinity tag at the N-terminus of PR was used in place of poly-histidine.^37, 42^

Strep-II is a minimal eight amino-acid peptide that binds to the core of streptavidin and has superior properties for efficient affinity purification of recombinant fusion proteins. Since the Strep-II tag is biologically inert and does not affect protein folding, it is potentially ideal for isolation of large multi-domain proteins like PR and CoRs in their intact native state.^43^ Strep II tagged PR-A or PR-B were expressed in Sf9 insect cells in the presence of the progestin ligand agonist R5020 to activate the receptor in culture. As described in more detail in Methods, receptors in native cell lysates were purified through two-step affinity and size-exclusion chromatography (SEC). The SEC fractions from the major protein peak that correspond with the expected molecular size of monomeric PR were pooled and concentrated in the range of 1.5mg-1.6mg/ml (15uM-18uM) with yields between 2.25 to 2.4mg of total protein from 1 liter of Sf9 cell cultures. Assessed by SDS-PAGE, each PR isoform contained a single major protein band of expected molecule size with a purity >98% (**Supplementary** Figure 1 **[Fig. S1]**). Yields were higher than with poly-histidine tagged PR from previous work, presumably due to the greater efficiency of Strep II tagged affinity system that results in fewer contaminating proteins with a single purification step. Mass spectrometry confirmed the identity of major protein bands of purified PR-A and PR-B as intact full-length PR-A or PR-B with no other peptides identified from unrelated insect cell protein background with a coverage >89% (**Fig. S2**). However, purified PR-B typically contained a faint band (∼2% of total) running slightly smaller than the major band, suggestive of a degradation fragment of PR. Further mass spectrometry analysis of the smaller sized band showed it is also intact full-length PR-B with a slightly different phosphorylation pattern than the major band (**Fig. S3**). Phosphorylation of PR is known to alter its mobility on SDS-PAGE.^36^

CoRs, SRC3 and p300, were expressed in Sf9 insect cells and purified in a similar two-step procedure as full-length constructs with Strep II tags at the amino terminus. This resulted in SRC3 (**Fig. S4)** and p300 **(Fig. S5**) exhibiting singular major bands on SDS-PAGE of expected molecular size at >98% purity. The concentrations of purified SRC3 and p300 were in the range of 2.0-2.5mg/ml (12uM and 9uM) with yields of 2.0-2.5 total mg from 1 liter of Sf9 cell cultures. Mass spectrometry confirmed the identity of SRC3 and p300 each as intact full-length protein with no detectable unrelated insect cell peptides (**Fig. S6**).

### Quality of purified proteins and DNA-induced dimerization of PR

The quality of purified proteins was further assessed by size-exclusion chromatography-multi-angle light scattering (SEC-MALS) and differential scanning fluorimetry (DSF). We performed SEC-MALS for each purified protein and DNA, alongside mixtures of the different proteins and DNA. SEC-MALS chromatograms for each protein displayed single peaks with experimentally derived molecular weights (MW) within the expected theoretical MW for each macromolecule as a monomer, except PR-B which behaved as a mixture of monomer and dimer **(Fig. 1 and Table S1).** Slight discrepancies between experimental and theoretical MW of individual proteins (e.g., PR-A, SRC3, p300) were likely due to deviation of dn/dc values from that of BSA. BSA is a globular protein while PR-A, SRC3 and p300 have significant disordered regions. In the absence of DNA, PR-A, bound to the progestin agonist ligand R5020, showed monomeric MW. Addition of DNA resulted in an experimentally determined MW within the expected theoretical for a PR-A dimer (**Fig. 1 and Table S1)**. PR-B liganded with R5020 gave a mixture of monomer and dimer MW distributions in the absence of DNA, and a single MW distribution within the expected theoretical for a PR-B dimer upon complex assembly with DNA **(Fig. 1 and Table S1).** These results are consistent with previous data showing that PR binds to PRE DNA as a dimer and further indicates that DNA induces formation of stable dimers. SEC-MALS also detected higher order complex assembly of all three proteins (PR-A R5020 agonist-bound, SRC3, p300) on DNA, eluting ahead of p300 analyzed alone **(Fig. S7**). The peak of the protein mixture had a long tail and showed heterogeneous MW distribution, indicating some overlap with monomeric p300 and SRC3. Experimentally determined MW analysis indicated formation of a 540 kDa complex for the peak apex, which could reflect weighted averages of a 1:1:2:1 p300:SRC3:PR-A:DNA complex (expected MW∼612 kDa) and monomeric p300 (expected MW=267 kDa). We observed the presence of DNA in the peak apex (**Fig. S7**) and subsequent disappearance of the SRC3 peak when all three proteins were added, confirming the formation of the higher order complex. These data are consistent with a stoichiometry of 1:1:2:1 (p300:SRC3:PR:DNA) in the complex, agreeing with the PR:SRC2:p300/DNA CryoEM structure previously reported.^19^

**Figure 1.**
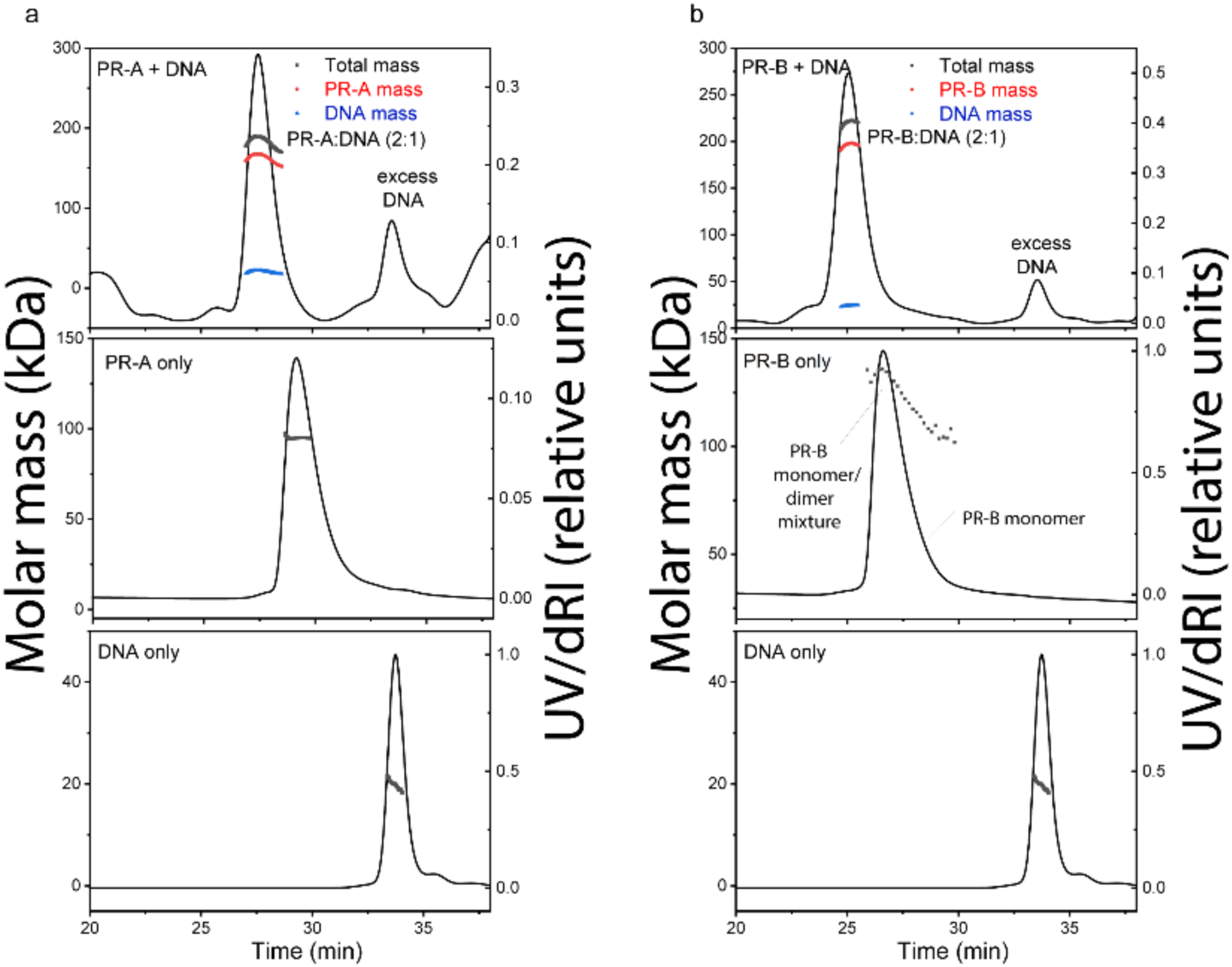
DNA induces assembly of PR-A and PR-B. ***A-B.*** SEC-MALS chromatograms of agonist (R5020)-bound PR-A (A) and PR-B (B) with and without DNA. The molar mass of DNA and PR-A alone matches the monomeric molar mass (black line/dots across the peaks). DNA induces assembly of both PR-A and PR-B into a complex with 2:1 (protein:DNA) stoichiometry. The presence of DNA in the complexes were confirmed by deconvolution of the protein and DNA fractions in the peak (red and blue lines, respectively).

DSF was used to determine the native folded state of each purified protein and stability to thermal denaturation. Proteins were incubated with SYPRO Orange, a dye that interacts with hydrophobic regions that become exposed during denaturation and can be used as a readout of protein unfolding. For all proteins analyzed (PR-A, PR-B, SRC3 and p300), thermal denaturation emission curves indicated cooperative unfolding transitions with melting temperatures (T_M_) in the range of 40°C to 49°C, indicative of moderately stable structured proteins **(Fig. S8, Table S2).** Addition of PRE DNA increased the T_M_ of PR-A and PR-B, consistent with increased stability of PR dimers as a complex with DNA **(Table S2).**

### DNA binding induces PR isoform specific conformational changes

Perturbations in deuterium exchange induced by DNA binding differ between the PR-A/PRE and PR-B/PRE complexes (**Fig. 2**). The intrinsic deuterium exchange of non-DNA-bound PR-A and PR-B showed similar exchange profiles between isoforms (**Fig. S9**), demonstrating that the differences in solvent exchange were not caused by inactive protein. Addition of PRE DNA resulted in decreased deuterium exchange (protection from solvent exchange) throughout the PR-A DBD and LBD, indicating DNA-mediated stabilization of these domains (**Fig. 2**). Differential HDX-MS of PR-B revealed fewer perturbations within regions of the NTD, DBD-CTE, and LBD in response to DNA than PR-A; however, both saw protections at the PR dimerization domains (PR-B amino acids: 602-618 and 885-922) through decreased deuterium exchange when DNA-bound (**Fig. 2**). These results agreed with those observed using SEC-MALS (**Fig S1, Table S1**). Thus, alterations in deuterium exchange are likely a combination of homodimerization and DNA binding. Interestingly, decreased solvent exchange was observed in the PR-B NTD (amino acids 270-276), which is not observed in PR-A, potentially indicative of a shift in NTD-LBD intraprotein interactions that only exist for PR-B.

**Figure 2.**
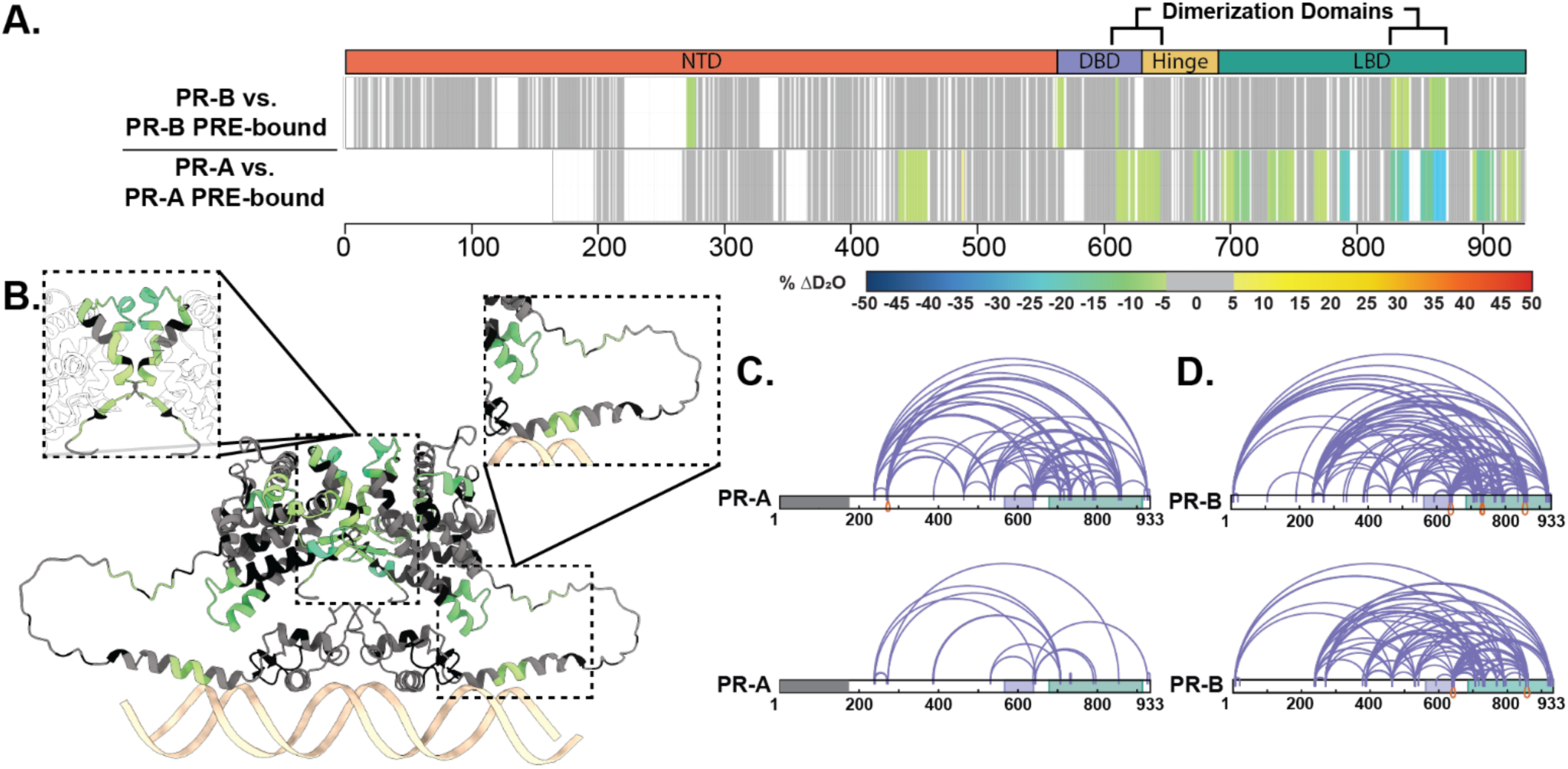
Structural proteomics reveals isoform differences upon response element binding. **A.** Consolidated HDX-MS data, run in triplicate, showing the differential analysis between unbound PR vs. PRE-bound where the top is PR-B, and the bottom is PR-A. Domains are labeled as the following: N-terminal Domain (NTD), DNA-binding domain (DBD), Hinge region (Hinge), and Ligand binding domain (LBD). **B.** Trimmed AlphaFold 3.0 model (residues 375-769) of PR-A homodimer with unbound PR-A vs. PR-A:PRE HDX overlays. Highlighted regions are the PR dimerization domain (top) and the DBD C-terminal extension (bottom). Cooler colors indicate stronger HDX protection. **C.** XlinkX images of differential PR-A ± PRE experiments, where crosslinks from the unbound (top) and PRE-bound (bottom) states are shown. Crosslinks mapped onto PR-B numbering with the gray region representing the 164 amino acids not expressed in PR-A. Results representative of triplicate experiments, with validation in Skyline. **D.** XlinkX view of differential PR-B ± PRE experiments, where crosslinks from unbound (top) and PRE-bound (bottom) states are shown. Domains – Gray: Not expressed; Purple: DBD; Green: LBD.

Crosslinking mass spectrometry (XL-MS) was used to assess amino acid proximity, which can inform inter- and intradomain interactions. XL-MS showed a rearrangement between the CTE and LBD when DNA was present, indicating compaction of those regions upon DNA binding (**Fig. 2**). Simultaneously, crosslinks from the PR-A N-terminus to C-terminus (residues 240 and 933) were diminished, indicating that these PR domains are no longer in proximity, most likely due to PR stabilization through homodimerization. PR-B crosslinks from the N-terminus to C-terminus (residue 7 to 933) were also lost upon DNA binding. The reduction of these N- to C-term crosslinks while others not near the C-term are retained, suggests a movement of the PR NTD away from the LBD. All together, these data show PR NTD stabilization through NTD-LBD interactions without DNA present. When bound to a canonical PRE, PR rearranges to shift the NTD slightly away from the LBD, and these interactions are not required for protein stability (**Fig. 2**).

### Co-regulator binding induces conformational changes to each PR isoform

Using a sequential CoR addition strategy, the differential deuterium exchange for PR was measured against the PR:SRC3 complex using the stoichiometries discerned from the SEC-MALS results (2:1:1; PR:SRC3:PRE; **Fig. S7**). Protections were observed throughout PR-A (**Fig. 3**). This meaning that SRC3 can bind PR-A as a monomer in the absence of DNA. SRC3-mediated exchange protection was not centralized around a specific PR-A domain, but the dimerization domains, NTD, and CTE all showed decreased deuterium exchange (**Fig. 3**).

**Figure 3.**
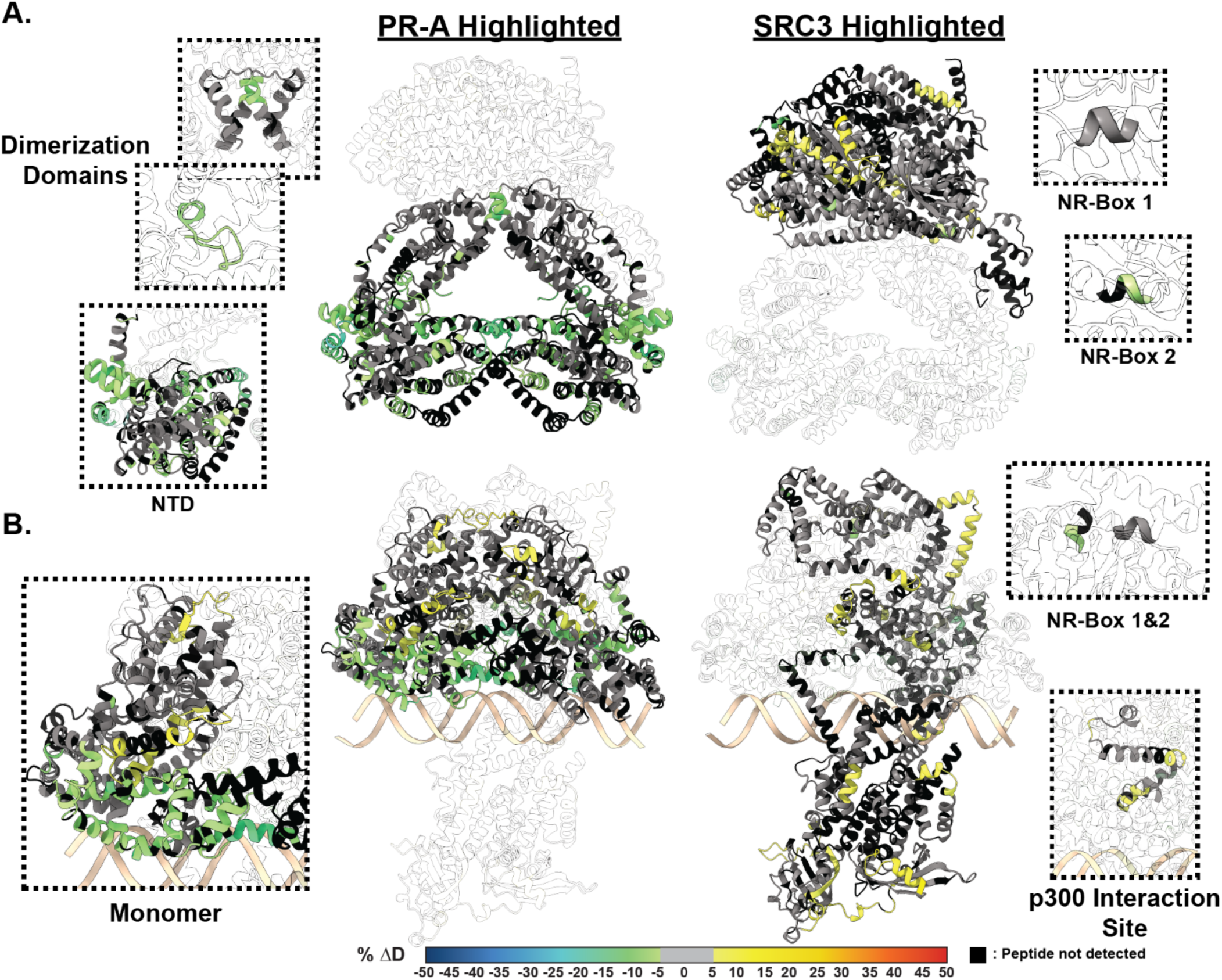
SRC3 induces LBD changes to PR upon PRE addition. **A. Left** HDX overlay (PR-A vs. PR-A:SRC3) mapped onto AlphaFold3.0 model of the PR-A:SRC3 ternary complex with the PR homodimer highlighted. Zoomed-in sections of PR corresponding to the dimerization domains (Amino acids: 720-769 and 438-454) and N-terminal domain (PR-A amino acids 1-476) highlighted with matching HDX overlays. **Right.** Differential HDX overlay of SRC3 vs. PR-A:SRC3 onto the best scoring PR:SRC3 apo complex with SRC3 highlighted. NR-Boxes 1 and 2 (amino acids 685-689 and 738-742, respectively) blown up to show differential exchange. **B. Left** HDX overlay (PR-A:PRE vs. PR-A:SRC3:PRE) mapped onto AlphFold3.0 model of PR-A:SRC3:PRE ternary complex with the PR homodimer highlighted. One PR-A monomer is shown as a zoomed-in section. **Right.** Differential HDX overlay of SRC3 vs. PR-A:SRC3:PRE onto the best scoring PR:SRC3 apo complex with SRC3 highlighted. NR-Boxes 1 and 2 and the p300 interaction site (amino acids 1023-1093) are highlighted to show differential exchange. Black peptide regions correspond to peptides not identified by HDX-MS. Each color represents the percent change in deuterium incorporation (Δ%D), following the scale shown at the bottom.

This was analogous to the previous ± DNA HDX-MS results (**Fig. 2**), indicating SRC3 may facilitate PR dimerization without requiring DNA. Further protections were observed for the PR-A NTD when DNA-bound, suggesting direct PR NTD and SRC3 interactions (**Fig. 3**). Interestingly, the PR-A LBD in the PR-A:SRC3 complex showed increased deuterium exchange (deprotection) when DNA-bound (**Fig. 3**. B). This was unexpected, as PRE addition was expected to enhance complex stability like that observed for the PR:PRE complexes (**Fig. 2**). Yet even though increased exchange was observed in the LBD, neither the dimerization domains nor AF-2 (H3, H4, and H12) were affected. This indicates that receptor binding either PRE, SRC3, or a combination of the two influences the conformational dynamics of the PR LBD without destabilizing the homodimer.

Similar exchange profiles were observed for PR-B in its SRC3-bound complexes (**Extended Data Fig. 1**). However, PR-B displayed fewer regions of differential exchange across the protein, compared to PR-A. In the PR-B:SRC3 complex, some protections were observed in the PR-B NTD; however, not throughout the rest of the protein (**Extended Data Fig. 1**). DNA binding does not seem to affect the PR-B:SRC3 complex, where few differences are measured between the PR-B homodimer and PR-B:SRC3 (**Extended Data Fig. 1**). Taken together, these data suggest a concerted SRC3-mediated deprotection of the PR LBD and protection of the DBD-CTE when DNA bound, and stabilization of PR when SRC3 bound in the absence of DNA. This may indicate an SRC3-mediated PR priming mechanism, where deprotections to the solvent boundary may increase the solvent-accessible surface area for additional CoR binding.

However, to understand how the HDX results translate to protein structure, AlphaFold3.0^44^ predictions were generated for the unbound and PRE-bound PR:SRC3 complexes. Since AlphaFold3.0 introduces spurious structural order (termed, hallucinations)^45^ to intrinsically disordered regions such as the NTD of PR, 25 independent models were generated by repeating structural predictions five times with random seed values (**Fig. S10**). These models were then examined with HDXer^46^ to identify the model that best fit the HDX-MS data. Each top-scoring model had average root mean square errors (RMSEs) less than 0.4.

The top model for non-DNA-bound and DNA-bound PR-A display differing SRC3 binding modalities, where SRC3 envelops the PR-A homodimer only in the DNA-bound model (**Fig. 3**). The non-DNA-bound model exhibits a distinct separation between PR and SRC3, showing the main interaction site as the AF2-cleft (**Fig. 3**). When bound to DNA, the homodimer binds to the DNA on one face while SRC3 energetically favors binding on the opposite side of the DNA (**Fig. 3**). Overlaying the HDX data, it was clear to see that the HDX-MS data indicate a simultaneous stabilization of the DBD-CTE and increased motility at helical connecting regions, shown via the HDX protections and deprotections (Fig. 3**.B, Left**). Representative PR-B:SRC3 AlphaFold models suggest a similar binding modality to PR-A. When bound to DNA, the best fit model has PR-B adopting an elongated structure (**Extended Data Fig. 1**). While the orientation of SRC3 is similar between each PR isoform, PR-B has enhanced protein-protein interactions due to its extended NTD (**Extended Data Fig. 1**).

XL-MS was then used to identify interprotein interactions. When non-DNA bound, PR-A and SRC3 have multiple interactions at the PR LBD (**Extended Data Fig. 2**). Adding the HDX to the crosslinking maps established that differential exchange was localized to inter-or intraprotein crosslinking regions, displaying the agreement between methods (**Extended Data Fig. 2**). These showed that the AF-2 protections are caused by SRC3 binding, and the deprotections aligned with PR-SRC3 interprotein crosslinks (**Extended Data Fig. 2**). Other regions across PR showed corroborating HDX-MS and XL-MS data, which gives greater confidence that the differential deuterium exchange was SRC3-mediated.

In a similar manner to the PR-SRC3 complex, p300 was added to the transcriptional complex to assess p300 interaction sites on PR using HDX-MS and XL-MS. When p300-bound, there was more than a 10% change in deuterium exchange for both PR and SRC3 (**Fig. 4**, **Fig. 5**). These protections show that p300 strengthens the ternary complex, whether through direct PR-p300 interactions or strengthening the PR-SRC3 interaction. This behavior was only measured with PR-A, where we found a concerted shift from deprotected to protected regions in the PRE-containing complexes (**Fig. 4**). This points to p300-mediated increased complex stability for the PR-A-containing complexes. Considering these effects were not measured for PR-B, it further suggests that complex formation is not dependent on direct PR-B-SRC3 interactions, but rather is p300-driven for PR-B.

**Figure 4.**
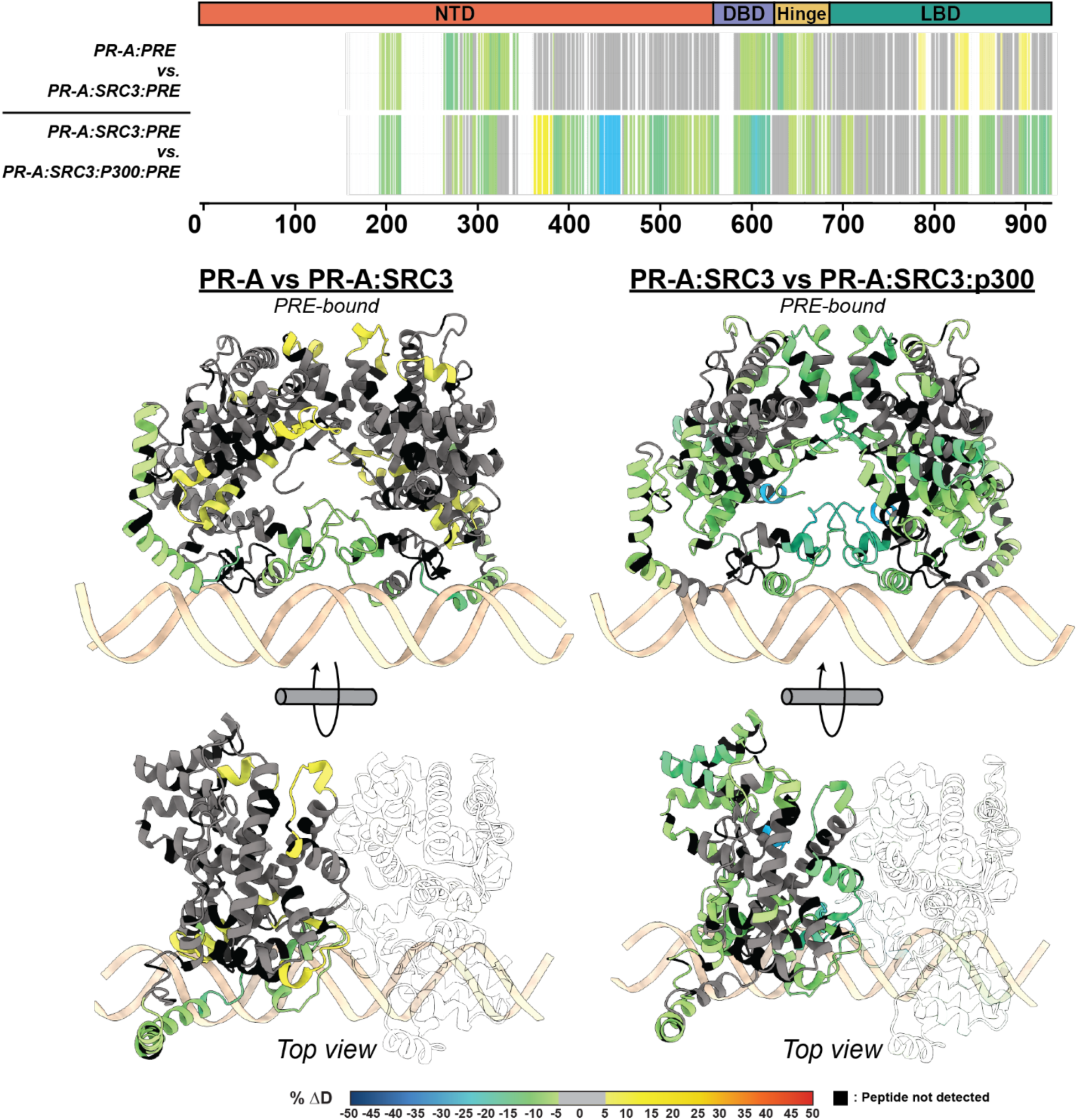
p300 differentially alters the conformational dynamics of PR-A and PR-B within the PR:SRC3:p300 complex. Top. Consolidated HDX plots of PR-B showing the differential HDX-MS comparisons within the plot to the left. Changes in deuterium uptake are represented by the rainbow plot shown at the bottom. Common PR domains are highlighted at the top: N-terminal domain (NTD, orange), DNA-binding domain (DBD, purple), Hinge (yellow), and ligand-binding domain (LBD, teal). **Bottom.** AlphaFold3.0 models of PR from the AF1 to LBD (amino acids 456-933 using PR-B numbering). HDX-MS overlays represent the same experiments as the consolidated views on the top. Each color represents the percent change in deuterium incorporation (Δ%D), following the scale shown at the bottom. Gray overlays indicate no significant changes and black indicates peptides not detected in the HDX-MS experiment.

**Figure 5.**
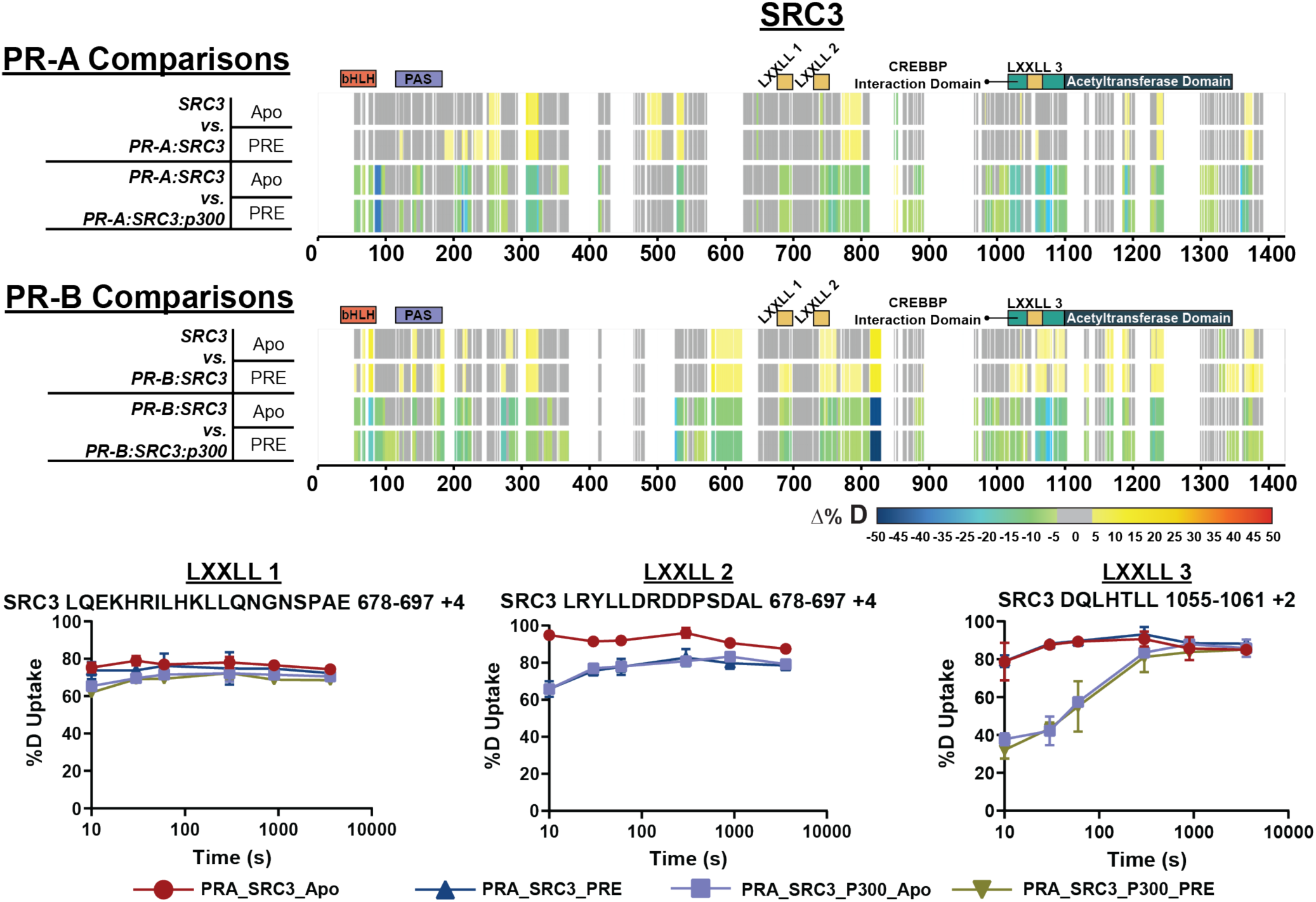
PR-A and PR-B differentially interact with SRC3 and are stabilized by p300 addition. **Top.** Consolidated differential HDX-MS results for SRC3, comparing the changes induced by PR-A and p300 binding in the presence and absence of PRE DNA. **Middle.** Consolidated HDX-MS plot of SRC3 exchange, with PR-B comparisons in the same order as PR-A. The motifs highlighted are the following: bHLH (orange), PAS (purple), LXXLL motifs (yellow), CREBBP Interaction domain (teal), and acetyltransferase domain (dark blue). Each color represents the percent change in deuterium incorporation (Δ%D), following the scale shown at the bottom.Gray overlays indicate no significant changes and black indicates peptides not detected in the HDX-MS experiment. **Bottom.** Selected deuterium uptake plots for peptides that contain LXXLL motifs 1, 2, and 3. The %D uptake indicates the percent deuterium uptake over time for the PR-A:SRC3 ± DNA and PR-A:SRC3:p300 ± DNA HDX experiments.

Crosslinking results trended similarly, where multiple interactions were found between PR, SRC3, and p300 (**Extended Data Fig. 3**). Multiple crosslinks across the three proteins point to increased stability of the PR-SRC3-p300 ternary complex, compared to the PR-SRC3 complex (**Extended Data Fig. 2**). Unlike from the HDX-MS data, XL-MS showed no major differences between isoforms. Each PR isoform showed a unique feature at the C-terminus, where the crosslinking between the three proteins converged. This coincided with the AF-2 domain of PR, acetyltransferase domain of SRC3, and NCOA2 interaction domain of p300, which could serve as a specific activation point for PR-mediated transcription.

### Structural proteomics identifies SRC3 and p300 NR-box utilization for PR isoforms

SRC3 contains conserved domains known as nuclear receptor (NR) boxes. NR-boxes are sequences that promote nuclear receptor binding with a motif of LXXLL. In the case of PR-SRC3 interactions, these indicate a potential PR binding site. For SRC3, there are three separate NR boxes where PR has the potential to bind (amino acids 685-689, 738-742, and 1057-1061). It is unknown, though, which or how many of these are necessary for the PR-SRC3 interaction. Prior data has identified PR utilization of a combination of NR-Box 1 and 2 for SRC1-mediated activation,^47–49^ yet this has not been established for SRC3. Using HDX-MS, deuterium exchange profiles can be monitored for multiple proteins within a singular experiment. Using the same results for assessing PR-specific exchange, solvent exchange differences were measured for SRC3 to identify PR and p300 binding sites. Differential HDX-MS experiments showed protections at NR-box 2 upon PR-A addition and non-significant (see: Methods – Data Rendering) decreases in NR-box 1 (**Fig. 5**). Interestingly at NR-box 3, increases in solvent exchange were observed, showing greater solvent accessibility at this region (**Fig. 5**). These data suggest that NR-box 2 is initially utilized by PR-A, NR-box 1 is utilized to a lesser extent, and the third NR-box remains unbound in both the PR-A apo and PRE-bound state.

Upon p300 addition, all NR-boxes were protected in the PR:SRC3:p300 and PR:SRC3:p300:PRE differential experiments, including NR-Box 3 (**Fig. 5**). Observing the percent deuterium uptake curves for representative peptides of each NR-Box, a nearly two-fold decrease in the deuterium exchange was measured for NR-Box 3 upon p300 addition (**Fig. 5**). Since NR-box 3 was not protected in the absence of p300 in the PR:SRC3 complex, this implies that p300 directly binds NR-box 3. This is further supported by the crosslinking data, where PR formed crosslinks near NR-boxes 1 and 2 of SRC3 and p300 crosslinks with the third NR-box in SRC3 (**Extended Data Fig. 3**). In addition, p300 induced widespread protection throughout PR and SRC3. Taken together, p300 stabilizes both PR and SRC3 in both the absence and presence of DNA.

Similar comparisons were made using PR-B, which showed unique deuterium exchange compared to PR-In the SRC3 ± PR-B experiments, no protections from solvent exchange were observed in any NR-Box. However, solvent exchange increased in NR-Boxes 2 and 3 upon PR binding, regardless of PRE presence. Increases to exchange in NR-Box 1 were only observed in the presence of PRE. Upon p300 binding, there was exchange protection in each NR-box, different than the sequential binding seen with PR-A (Figure 5). The other SRC3 domains, though, have a similar protection pattern to PR-A where the bHLH, PAS, CREBBP interaction domain, and putative acetyltransferase domains were protected only in the presence of p300, most likely indicating direct p300 binding at those regions. This also supports that PR-A is the primary binding protein of SRC3 instead of PR-B, due to the protections seen at each NR-Box. The crosslinking showed that PR-B-SRC3 interactions were within NR-boxes 1 and 2, whereas NR-box 3 only contains crosslinks between SRC3 and p300, which was the same as PR-A (**Extended Data Fig. 3**).

To gauge the necessity of each NR-box for PR transcription, promoter:reporter assays were used in conjunction with SRC3 site-directed mutagenesis to measure PR transcriptional output. PR transcriptional response was indirectly measured using a PRE-firefly luciferase reporter plasmid.^50^ Mutating SRC3 NR-Box 2 from LXXLL to LXXAA resulted in reduced PR response (**Extended Data Fig. 4**). Changing other NR-Box sequences to LXXAA at either Box 1 or Box 3 did not result in reduced PR activity (**Extended Data Fig. 4**). However, combination mutants: NR-boxes 1+2 and 1+3 did reduce PR activity. The SRC3 Box 2+3 mutant increased overall activity (**Extended Data Fig. 4**). Surprisingly, the mutation of all three NR-boxes did not show any significant differences between WT PR-B and WT-SRC3, suggesting a potential compensatory mechanism by the other p160 coactivators. These results show NR-box 2 is an important PR interaction site for transcriptional response, and combination mutants involving NR-Box 1 are deleterious to PR activity. These assays were repeated with PR-A, but the results were inconclusive due to intrinsically reduced transcriptional response (**Extended Data Fig. 4**).

Previous data had shown that sequential addition of p160 CoRs and p300 influences NR transcriptional response.^51, 52^ However, the crosslinking results of the PR:SRC3:p300 ternary complex suggested direct PR-p300 binding (**Extended Data Fig. 3**). Based on these crosslinking results, direct PR-p300 interactions were investigated by XL-MS. In the absence of SRC3, PR and p300 formed crosslinks, indicating direct protein-protein interactions (**Fig. 6**). Crosslinks to both the N-terminus and C-terminus show that there are multi-pronged PR-p300 interactions without SRC3. These included crosslinks to the p300 NR-boxes, bromodomain, and core acetyltransferase regions. PR transcriptional response was also augmented when both PR and p300 were overexpressed, without SRC3 overexpression (data not shown).

**Figure 6.**
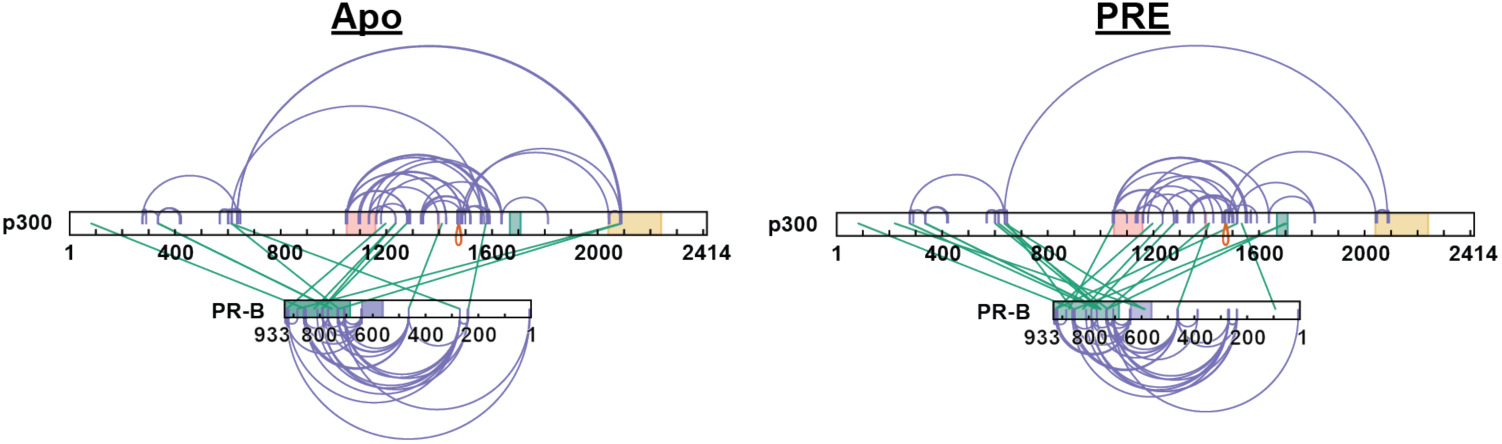
XL-MS shows p300 directly interacts with PR. **A.** Crosslinking results from PR-B:p300±PRE experiments. Purple: intraprotein crosslinks, green: interprotein crosslinks. PR highlighted domains: DBD (purple) and LBD (green). p300 highlighted domains: bromodomain (pink), zinc finger domain (green), and NCOA2-interaction domain (yellow).

### RU486-antagonism reorganizes PR:SRC3 and PR:SRC3:p300 protein complexes

The logical sequitur for R5020-bound protein led us to investigate the organization of PR complexes, bound to the progestin antagonist, RU486. As determined by SEC-MALS, both PR-A and PR-B bound to RU486 showed monomeric MW distribution in the absence of DNA (**Extended Data Fig. 5**). With addition of DNA, both PR isoforms assembled as a DNA complex with an experimentally determined MW within the expected theoretical for PR-A and PR-B dimers **(Extended Data Fig. 5, Table S1).** This behavior of purified PR bound to RU486 is consistent with previous studies demonstrating RU486 promotes efficient PR dimerization and binding to PRE DNA.^3, 53, 54^ Prior work with PR bound to RU486 showed that the C-terminal tail of the LBD adopts a distinct conformation from that of agonist bound PR^55, 56^ and X-ray crystallography of PR LBD bound to RU486 results in a displacement of helix 12 in multiple potential conformations that may interfere with formation of an AF-2 interaction surface for binding LXXLL motifs of Co-activators.^21^ X-ray crystallography of the PR LBD bound to antagonists related to RU486 further show a displacement of helix 12 from agonist conformation and differential binding of a peptide from the corepressor SMRT.^57^ Studies to date have not explored the influence of RU486 on interactions of PR and CoRs, each as full length proteins to explore alterations in interaction surfaces outside of the LBD.

We utilized a similar strategy as the R5020-bound PR experiments to define the PR-CoR relationship in the inactive conformation. XL-MS and HDX-MS showed different PR-SRC3 binding modalities than agonist-bound PR. In the agonist-bound complex, PR-SRC3 crosslinks group near the C-terminus of both proteins. However, when RU486-bound, the enriched crosslinks shift from a C-terminal to an N-terminal grouping (**Fig. 7**). This suggests that the proteins can still interact, even in an ‘inactive’ state. Further crosslinking in the PR-SRC3-p300 ternary complex showed crosslinks between PR, SRC3, and p300, suggesting all three proteins can interact when antagonist-bound. The most apparent difference for the RU486 crosslinking was the concerted loss of crosslinking between PR and p300 at the C-terminus (**Fig. 7**). This may be indicative of a transition from an active to inactive state for PR:SRC3:p300 driven transcription. Moreover, the crosslinks between PR and NR-Boxes 1 and 2 within SRC3 were lost upon RU486 binding of the receptor. This supports that the protein is in its inactive state yet can still form contacts with SRC3 and p300. This challenges the classical NR model for activation, where antagonist-bound protein cannot interact with coactivators.

**Figure 7.**
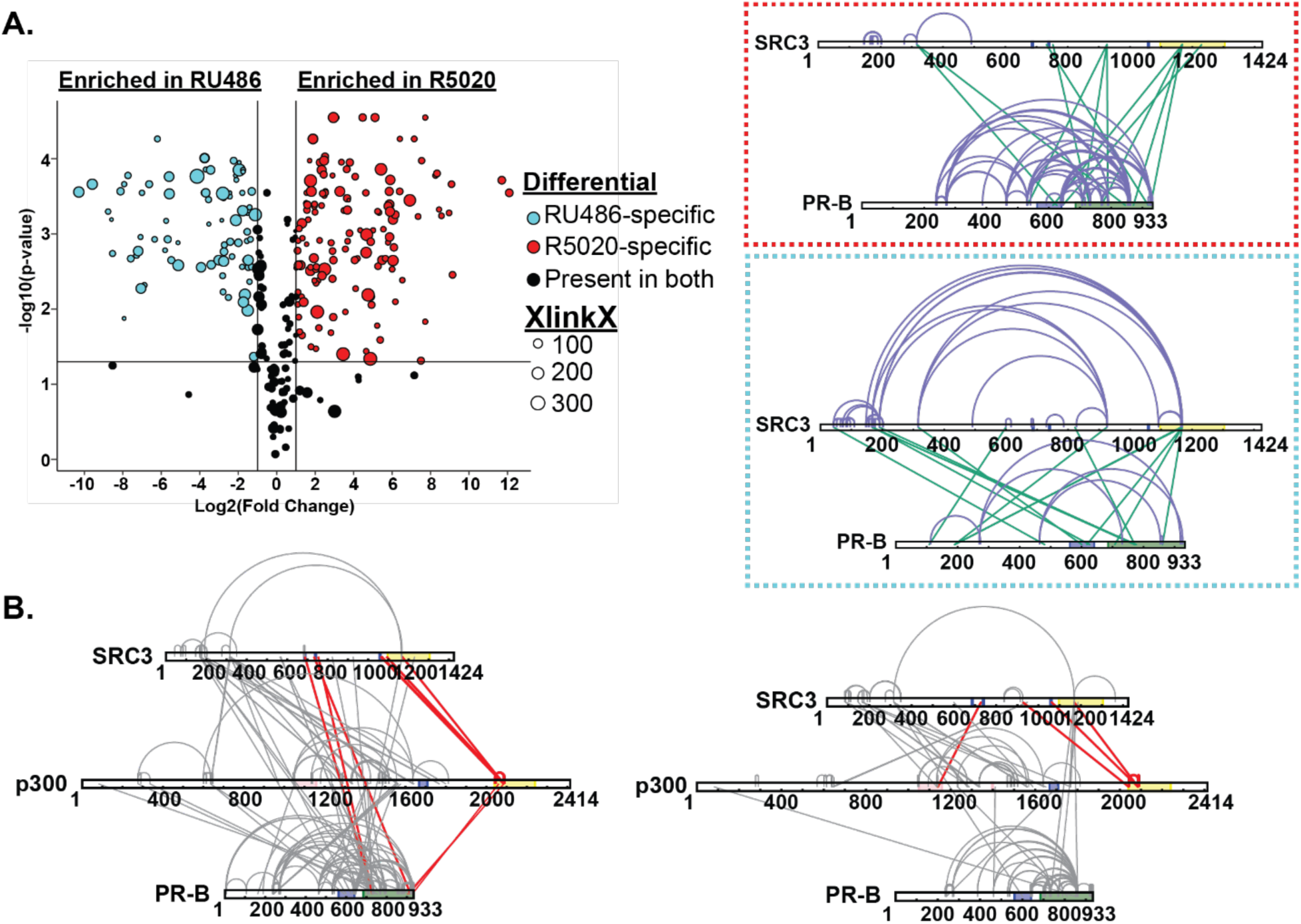
RU486 antagonism rearranges PR-SRC3-p300 interactions. **Top.** Plotted differential crosslinks in PR:SRC3 experiments, comparing R5020-specific (agonist, red) and RU-486-specific (antagonist, blue) crosslinks. Crosslinks represented by circles with corresponding XlinkX scores as point size. **Top. Red.** XlinkX view of R5020-specific crosslinks in differential PR:SRC3 experiments. **Top. Blue.** XlinkX view of RU486-specific crosslinks in differential PR:SRC3 experiments. **B.** All validated R5020-bound (left) and RU486-bound (right) crosslinks for differential PR-B:SRC3:p300 ± PRE experiments. The x-axis represents the Log2 transformed fold change values from Skyline, while the y-axis represents the -log10 transformation of the Skyline p-value output. The lines are indicative of a Log2 fold change of 1 (two-fold increase) and -log10 p-value of 1.3, corresponding to p<0.05. Crosslinks highlighted in red show notable differences in PR:SRC3:p300 interactions between PR ligands. Defined domains are as follows: PR - DBD (purple) and LBD (green); SRC3 – NR-boxes (purple) and histone acetyltransferase domain (yellow); p300 - bromodomain (pink), zinc finger domain (green), and NCOA2-interaction domain (yellow).

HDX-MS was applied to investigate the backbone dynamics of each protein. Distinctive from the R5020-bound complexes, the presence of antagonist significantly reduced deuterium exchange, specifically in all the PR ± PRE experiments, leading to reduced deuterium exchange throughout the protein (**Fig. S11**). However, the sequential addition of co-regulators showed an expected solvent exchange profile, comparatively. Because on this, comparisons were made exclusively between the larger ternary complexes. Interestingly, regions of protection were observed for the larger ternary complexes with both PR-A and PR-B for ± DNA experiments (**Fig. 8**). The differential deuterium uptake was similar to that seen with agonist (R5020)-bound PR in **Fig. 3**. Protected regions were localized to the DBD, CTE, and LBD, all regions known to interact with DNA and CoRs. In addition, portions of the AF-1 and AF-2 cleft had reduced deuterium exchange (**Fig. 8**), which is a hallmark of CoR binding. The most notable result was that CoR addition resulted in a stronger stabilization of both PR dimerization domains. This suggests that when antagonist-bound, PR can still associate with co-activators: SRC3 and p300, and these CoRs still help to stabilize PR in a complex.

**Figure 8.**
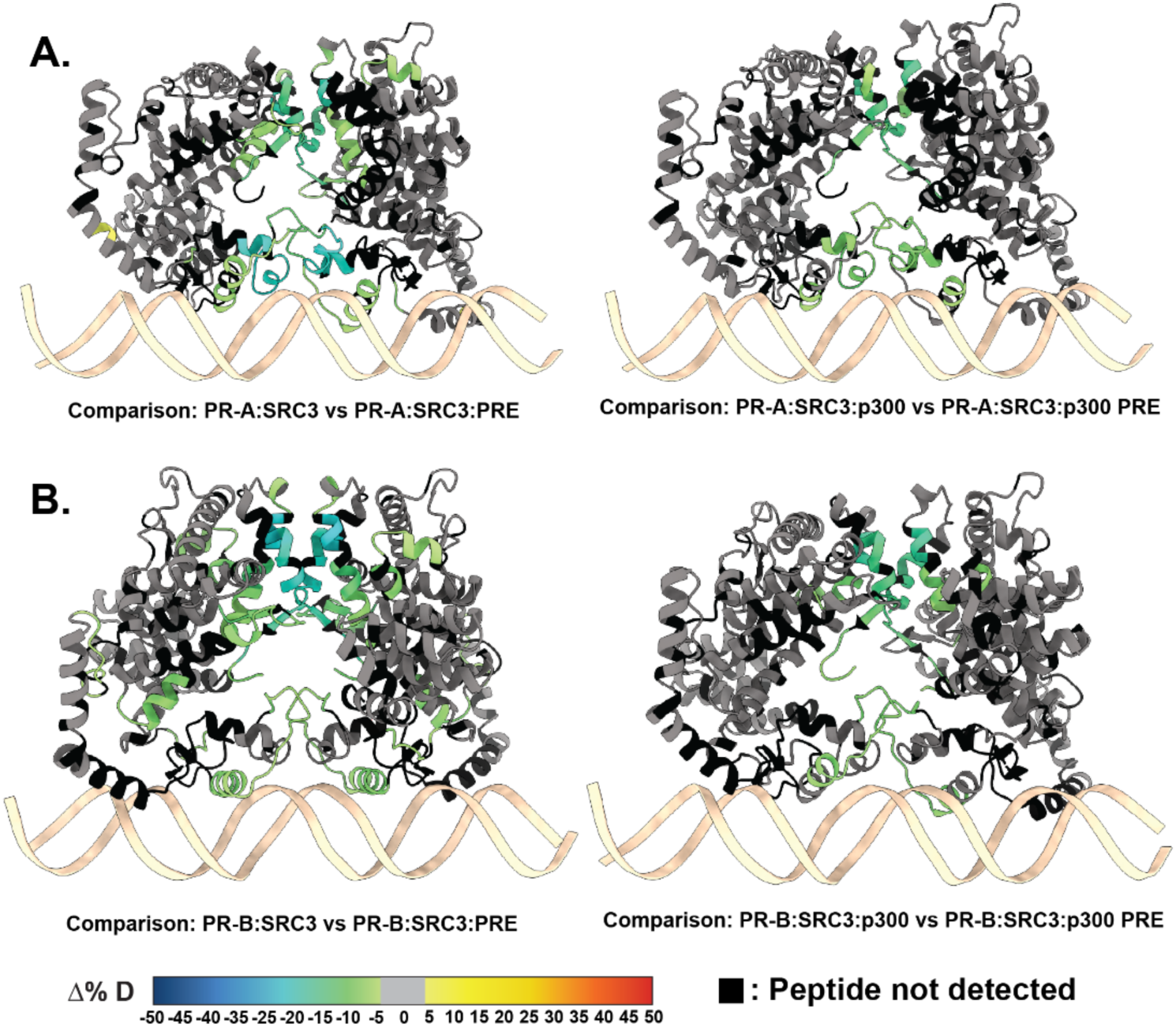
RU486-bound PR has reduced deuterium exchange upon CoR binding. **A.** PR models of AF1 to C-terminus (amino acids 456-933) with PR-A HDX overlays, corresponding to the comparisons shown beneath them. A. PR models of the AF1 to C-terminus with corresponding PR-B HDX overlays labeled beneath. Each color represents the percent change in deuterium incorporation (Δ%D), following the scale shown at the bottom. Gray overlays indicate no significant changes and black indicates peptides not detected in the HDX-MS experiment.

## Discussion

Steroid nuclear receptors (SR) are hormone-responsive transcription factors that exhibit remarkable functional diversity in mediating cell and target gene specific responses. These responses are largely driven by conformational dynamics of the receptor, enabling their binding of unique subsets of transcriptional CoRs and DNA response elements. Because of their presence in various cancer phenotypes, SRs have been attractive targets for cancer therapeutics through disruption of their transcriptional activity. In this work, PR was investigated to better understand the mechanism of the PR-CoR relationship. Each PR isoform, PR-A and PR-B, exhibits distinct physiological roles dependent on the cell or tissue type. The mechanistic basis for differences between isoform activity was not well defined. One recent low-resolution cryoEM structure of PR-B gave some clues about how such complexes may be formed; however, the lack of atomic-level structural details limited the understanding of interactions between PR-B and the CoRs used in that study. Moreover, structural understanding of PR-A and PR-A:CoR complexes was completely lacking. One of the associated problems in solving high-resolution structural analyses has been generation of stable PR:CoR complexes. A potential solution to this problem is using CoR peptides specific to regions of interest, but that excludes the interplay between protein secondary structures within a complex. Here, we make technical developments by improving protein coverage for structural proteomics studies, utilizing full-length PR and CoRs. Our studies clearly demonstrate generation of stable protein complexes with expected stoichiometry. This paved the way not only for carrying out structural dynamic studies of PR isoforms and their interactions with CoRs with or without bound to DNA but also for future structural studies for which finding a stable complex has been challenging.

Here, XL-MS and HDX-MS were used to make solution-phase structural measurements of amino acid distance constraints between protein components and to probe conformational dynamics of PR-A and PR-B in complex with DNA and CoRs. As expected, our results show both PR-A and PR-B binding to DNA as dimer. It had been hypothesized that DNA binding induces compact structure particularly in the intrinsically disordered NTD and other structurally flexible regions. The differential XL-MS data showed DNA-mediated changes to PR organization. Crosslinks enriched in DNA-bound PR indicated that DNA induces LBD movement away from the NTD, suggesting that PR converts from a compact to elongated structure upon DNA binding. These are interesting findings and provide much needed information regarding structural dynamics of PR and would likely apply to other members of the SR sub-family.

The HDX-MS experiments were imperative for us understanding the role each CoR plays in PR transcriptional complex formation. Reduced deuterium exchange observed throughout PR-A following the binding of SRC-3 – particularly in the PR dimerization region – suggested that that SRC3 may induce PR-A dimerization without the addition of PRE DNA. This mimics what would happen in a biological system, where PR is stabilized by SRC3 before binding to genomic DNA after release by heat shock proteins. Once DNA bound, we observed increases in deuterium incorporation that were unexpected, but these increases were not in PR dimerization or AF-2 domains. Based on these observations, it can be hypothesized that such conformational dynamic changes may be important in preparing PR-A surfaces for interactions with other CoRs and facilitate the process of the exclusion or inclusion of specific CoRs in the complex. This is a phenomenon commonly adopted by SRs, where distinct CoRs bind SRs when moving from one DNA site to another one in the genome.^58–60^ However, the differential deuterium exchange was not as pronounced in PR-B, suggesting PR-A is the primary interacting isoform in the PR-SRC3 ternary complex. This differs from prior work showing SRC1 and SRC2 preference for PR-B in other cell types.^61–65^ SRC3 is known to be a pathogenic CoR in breast cancer. The preference of SRC3 to bind PR-A gives us reason to believe this interaction could be a primary cause for why excess PR-A leads to worse outcomes in breast cancer, considering SRC3 is an independent cancer driver currently being targeted for therapeutics.^66^

The perturbations in differential PR HDX, induced by SRC3 and p300, showed that these CoRs act in a mutually beneficial fashion. This was evidenced by the initial deprotected regions within PR when SRC3 bound, which became protected once p300 was added. Using this sequential addition strategy, we assessed the contribution of each CoR on PR deuterium exchange to show that SRC3 prepares PR for additional CoR binding while p300 stabilizes the ternary complex. Aside from the PR changes, we identified NR-Boxes 1 and 2 as being the main PR interaction sites on SRC3, while p300 exclusively used NR-Box 3, confirmed by both HDX and XL-MS. These findings indicate a probable transcriptional activation mechanism in PR-SRC3-p300-driven transcription, where the C-terminal interactions serve as an activation trigger.

Lastly, we found that RU486 (antagonist) bound PR was still capable of binding SRC3 and p300. This challenges the classical nuclear receptor activation model, whereby nuclear receptors cannot interact with activating CoRs when SRs are antagonist bound. This finding leads us to believe that activation mechanisms are not as simplistic as previously thought. There is evidence that interaction surfaces change between the proteins to prefer an activation-based or repression-based transcriptional response depending on the ligand. However, cognate CoRs may still transiently interact with these proteins.

Together, enhanced expression and purification methods produce more stable, purer protein than previously described, which enabled a detailed mechanistic study on the PR transcriptional complex. Structural proteomics, through HDX and XL-MS, bridged the gap between a passive understanding of PR-CoR organization to provide more detail than previously reported. AlphaFold3.0-driven structure predictions indicated that PR:SRC3 complexes have unique structures depending on their DNA occupancy, and the experimental results presented reveal a sequential priming mechanism for PR. P300 further interacts with both the N- and C-termini of PR, positioning its HAT domain in close proximity to DNA and chromatin. Additionally, NR box preference indicates a specific order of complex assembly. Surprisingly, and contrary to classical models, SRC3 and p300 remain associated with PR when bound to antagonist, although the orientation of the complex differs from that observed with an agonist suggesting antagonist driven rearrangement of the transcriptional complex. Collectively, the findings presented here elucidate how SRC3 and p300 interact at the amino-acid level with PR-A and PR-B, influencing its function and conformation.

## Methods

### Materials

Oligonucleotides (Sigma-Aldrich). 32-mer progesterone response element (PRE) dsDNA sequences: Sense: 5’-CATCTTTGAGAACAAACTGTTCTTAAAACGAG-3’; Antisense: 5’-CTCGTTTTAAGAACAGTTTGTTCTCAAAGATG-3’. Sigma-Aldrich provided Disodium hydrogen phosphate (Na_2_HPO_4)_ (S9763), Sodium Chloride (NaCl) (S9888), Urea (51456), Benzonase (9025-65-4) and 2-Mercaptoethanol (2ME) (M3148). Fisher Scientific provided Glycerol (BP229-4), and Biotin (B0463). Hampton Research provided TCEP Hydrochloride (50-093-0). Invitrogen provided UltraPure™ 0.5M EDTA (15575020) and SYPRO Gel filtration standard (S6650). BIO-RAD provided unstained protein standard (1610363) and Gel Filtration Standard (1511901). Thermo Fisher provided Zeba™ Spin Desalting Columns 7K MWCO 0.5 mL (89882) and Disposable PES Bottle 0.2uM Top Filters (166-0045).

### Plasmids

Progesterone receptor protein-expression plasmids: pRP[Exp]-EGFP/PuroEF1A>hRluc(ns):P2A:hPGR[NM_000926.4] (co)* and pRP[Exp]-EGFP/PuroEF1A>hRluc(ns):P2A:hPGR[NM_001202474.3] (co)*, were constructed using VectorBuilder. Their respective vector IDs are VB230919-1497uvg and VB230919-1498ukn, which can be used to retrieve detailed information about the vector at vectorbuilder.com. Progesterone response plasmid, 4X PRE TK luc, was a gift from Renee van Amerongen^50^ (Addgene plasmid # 206159 ; http://n2t.net/addgene:206159 ; RRID:Addgene_206159).

### Protein expression

Human PR-A, PR-B, SRC3, and p300 as full-length open reading frame DNA were each synthesized with optimal codon usage for insect cells with an in-frame Strep II tag (WSHPQFEK/G) and a glycine spacer at the amino-terminus and inserted into pFastBac1 transfer vectors (Epoch Life Sciences, Houston, TX).

Recombinant bacmids were generated and expanded in Spodoptera frugiperda (Sf9) cell cultures and viral titers were determined by plaque assays as previously described.^38, 39^ Multiple 500 mL cultures of SF9 cells were infected with recombinant virus at an MOI of 2.0 and incubated for 48 hour at 27°C in oxygenated spinner vessels. For cells expressing PR-A or PR-B, the progestin agonist R5020 or antagonist RU486 was added to Sf9 cell cultures at 250 nM for 24 hours post-infection. Cells were collected and centrifuged 1500x g for 10min and pellets were washed once PBS by resuspension and centrifugation. Cell pellets were flash frozen and stored at -80°C.

### Protein purification

Cells (from 2x 500ml cultures) were resuspended in 50 mL of lysis buffer (50 mM Na_2_HPO_4_, pH 8.0, 500 mM NaCl, 5% Glycerol, 1 M Urea, 1 mM EDTA, 1 mM 2ME), supplemented with protease inhibitor tablets (leupetin, aprotinin, bacitracin, and PMSF) and submitted to Teflon-glass homogenization for 8 strokes at 1.5 speed in the cold room (4°C). The homogenate was treated with 1.2 units/mL of benzonase nuclease or 1 hour at 4°C then passed 3 times through an 18G needle followed by a 25G needle. The lysate was centrifuged twice for 1 hour at 50,00 xg and the resulting supernatant was filtered using 0.2 μm disposable filters. Purification was performed on AKTA Pure^TM^ 25 at 4 °C while monitoring conductivity and UV. A 1 mL StrepTrap XT prepacked Hi-Trap chromatography column (Cytiva) was equilibrated with lysis buffer for 10 column volumes (CV), and after loading the cleared cell lysate, the column was washed for 30 CV with equilibration buffer and then eluted with equilibration buffer containing 40 mM biotin. Eluted fractions (1 mL) were collected and analyzed by Coomassie Blue-stained 4-15% gradient SDS-PAGE gel. Fractions containing proteins of interest were pooled and concentrated to 1-2 mL by an Amicon ultracentrifugal device with a 10,000 MW cutoff (A280<4.0). Proteins were further purified by a preparative size exclusion chromatography (SEC) column (HiLoad 16/600 Superdex 200pg), degassed and equilibrated in SEC buffer (20 mM Hepes, pH 7.5, 200 mM NaCl, 5% Glycerol, 1 M Urea and 1 mM TCEP). Elution fractions (1 mL) were collected and analyzed by Coomassie Blue-stained 4-15% gradient SDS-PAGE gels and peak fractions were concentrated as above. Concentrations of purified protein products were by determined by Nanodrop absorbance at 280/260 nm, calculation of extinction coefficient, and by comparison of purified bands on Coomassie Blue-stained 4-15% SDS-PAGE gel with a standard curve of know amounts of unstained protein markers. Aliquots (50-100 μL) of purified protein were snap frozen in liquid nitrogen and stored at -80°C for up to 3-4 months and samples were used only once after thawing.

### Analytical SEC

All procedures were conducted at a temperature range of 0°C to 4°C.. Protein samples (50 µL each) were injected using a capillary syringe into a 50 µL loop and fractionated over a Superdex 6 Increase 5/150 GL column. The column was equilibrated with degassed SEC Buffer (20 mM Hepes, pH 7.5, 200 mM NaCl, 5% Glycerol, 1 M Urea, and 1 mM TCEP) as well as degassed SEC buffer without Urea (20 mM Hepes, pH 7.5, 200 mM NaCl, 5% Glycerol, and 1 mM TCEP). Columns were calibrated using Gel Filtration Standards.

### Differential Scanning Fluorimetry (DSF)

The thermal stability of SII-tagged purified proteins were measured by differential scanning fluorimetry (DSF) using the CFX96 Touch Real-Time PCR Detection System (BIO-RAD) instrument. Fluorescence monitoring was performed at 10°C–95°C at a rate of 0.5°C/s. Each reaction contained 1 μL 500x SYPRO^TM^ Orange gel stain, 4 μg purified protein and storage buffer which brought the reaction volume to 20 μL. All samples were scanned 3 times as technical replicates. Melting temperatures (T_m_) were estimated using the Bio-Rad CFX Maestro computer program by fitting the Boltzmann sigmoidal curve to the normalized data, and values represent an average of 3 replicates plotted with standard error of the mean.

### Size Exclusion Chromatography-Multi-Angle Light Scattering (SEC-MALS)

Data were collected using a Dawn Ambient light scattering instrument equipped with a 661 nm laser (Wyatt). The whole system is linked to an HPLC system with UV absorbance detection at 280 nm (Agilent) and an Optilab (Wyatt) for differential refractive index (dRI) measurements. Approximately 20 to 100 µg of proteins (p300, SRC3, PR-A (agonist R5020 or antagonist RU486), PR-B (R5020 or RU486) or DNA alone or in complexes were injected and flowed through a Superose 6 increase column (Cytiva). Data was analyzed using the Astra software (Wyatt). BSA sample was also run as a calibration control and to obtain the dn/dc values in the different buffer conditions. The buffer was 20 mM Hepes, 50mM NaCl, 5% glycerol, 1 mM TCEP, with or without 1 M Urea, pH 7.5. Figures were plotted using the Origin software. Standard errors shown were 5% of the calculated MW.

### Hydrogen-Deuterium Exchange Mass Spectrometry

#### Peptide identification

Peptides were identified using MS/MS experiments performed on a QExactive (ThermoFisher Scientific, San Jose, CA) over a 70-min gradient. Product ion spectra were acquired in data-dependent mode, and the five most abundant ions were selected for the product ion analysis per scan event. For peptide identification, the MS/MS *.raw data files were analyzed on Sequest (version 2.3, Matrix Science, London, UK). Mass tolerances were set to ± 0.6 Da for precursor ions and ± 10 ppm for fragment ions. Oxidation to methionine was selected for variable modification. Non-specific digestion was selected in the search parameters with 4 missed cleavages. Only peptides with an FDR<1% were used in the dataset, and duplicate charge states were used for each peptide in the peptide set.

#### Continuous labeling

Experiments with continuous labeling were carried out on a fully automated system (CTC HTS PAL, LEAP Technologies, Carrboro, NC; housed inside a 4°C cabinet) as previously described^28^ with the following modifications: For differential HDX, protein-protein complexes were generated by sequential protein addition, where each protein would incubate for 15 minutes at 4°C before sequential addition of the next protein. For larger complexes the order was as follows: DNA, PR, SRC3, then p300, waiting 15 minutes between each addition. The final incubation was carried out in the sample plate for 30 minutes before the experiment started (1-hour total incubation time from final protein addition). The reactions (5 μL) were mixed with 20 μL of deuterated (D2O-containing) buffer [20 mM Hepes, 200 mM NaCl, 1M Urea, 1 mM TCEP, and 5% glycerol (pD 7.9)] and incubated at 4°C for 0, 10, 30, 60, 900, or 3600 s. Following on-exchange, unwanted forward- or back-exchange was minimized, and the protein complex deuteration was stopped by the addition of 25 μL of a quench solution [20 mM NaH_2_PO_4_ and 1% TFA (pH 2.5)] before immediate online digestion and data acquisition.

#### HDX-MS analysis

Samples were digested through an immobilized fungal XIII/pepsin column (1-to-1 ratio, prepared in-house) at 50 μL/min [0.1% (v/v) TFA at 4°C]. The resulting peptides were trapped and desalted on a 2 mm-by-10 mm C8 trap column (Hypersil Gold, ThermoFisher Scientific). The bound peptides were then gradient-eluted [4 to 40% (v/v) CH_3_CN and 0.3% (v/v) formic acid] on a 2.1 mm-by-50 mm C18 separation column (Hypersil Gold, ThermoFisher Scientific) for 5 min. Sample handling and peptide separation were conducted at 4°C. The eluted peptides were then ionized directly using electrospray ionization, coupled to a high-resolution Orbitrap mass spectrometer (QExactive, ThermoFisher Scientific).

#### Data rendering

The intensity-weighted mean m/z centroid value of each peptide envelope was calculated and converted into a percentage of deuterium incorporation. This is calculated by determining the observed averages of the undeuterated (t=0 s) and fully deuterated spectra using the conventional formula described elsewhere.^26^ The fully deuterated control, 100% deuterium incorporation, was calculated theoretically, and corrections for back-exchange were estimated to be 70% deuterium recovery, accounting for 80% final deuterium concentration in the sample (1:5 dilution in deuterated buffer). Statistical significance for the differential HDX data is determined by an unpaired t-test for each time point, a procedure that is integrated into the HDX Workbench software.^67^

The HDX data from all overlapping peptides were consolidated to individual amino acid values using a residue averaging approach. For each residue, the deuterium incorporation values and peptide lengths from all overlapping peptides were assembled. A weighting function weights shorter peptides more than longer peptides. Weighted deuterium incorporation values were then averaged to produce a single value for each amino acid. The initial two residues of each peptide, as well as prolines, were omitted from the calculations. This approach is similar to one previously described.^68^

Deuterium uptake for each peptide is calculated as the average of %D for all on-exchange time points, and the difference in average %D values between the unbound and bound samples is presented as a heatmap with a color code given at the bottom of each figure (warm colors for deprotection and cool colors for protection). Peptides are colored by the software automatically to display significant differences, determined either by a greater than 5% difference (less or more protection) in average deuterium uptake between the two states or by using the results of unpaired t-tests at each time point (P < 0.05 for any two time points or P < 0.01 for any single time point). Peptides with nonsignificant changes between the two states are colored gray. The exchange of the first two residues for any given peptide is not colored. Each peptide bar in the heatmap view displays the average Δ %D values, associated SD, and the charge state. In addition, overlapping peptides with a similar protection trend covering the same region are used to rule out data ambiguity. Outputs from data rendering were transferred to ChimeraX (version 1.8) for generating data overlays onto predicted models.

These data have been deposited to the ProteomeXchange Consortium via the PRIDE^69^ partner repository with the dataset identifier PXD056400 for continuous labeling HDX-MS experiments.

### Crosslinking Mass Spectrometry

#### Sample preparation

For DSSO (ThermoFisher, A33545) cross-linking reactions, PR was diluted to 5 μM in cross-linking buffer [20 mM Hepes (pH 7.5), 200 mM NaCl, 1 M Urea, 1 mM TCEP, and 5% glycerol] and incubated for 30 min at room temperature before initiating the cross-linking reaction. For multi-protein complexes, DNA was added, then PR, then SRC3, then p300, waiting 15 minutes between each sequential addition. The proteins were added at consistent molar ratios (PR:SRC3:p300:DNA; 2:1:1:1.5) as determined in this paper, scaling each to PR at 5 μM concentration, total maximum protein concentration equal to 10 μM. The final incubation was performed for 30 minutes at room temperature. DSSO cross-linker was freshly dissolved in the cross-linking buffer to a final concentration of 150 mM before being added to the protein solution at a final concentration of 3 mM. The reaction was incubated at 25°C for 45 min and then quenched by adding 1 μL of 2.0 M tris (pH 8.0) and incubating for an additional 10 min at 25°C. Control reactions were performed in parallel by adding DMSO instead of crosslinking reagent. All cross-linking reactions were carried out in three replicates. The presence of cross-linked proteins was confirmed by comparing to the no-crosslink negative control samples using SDS-PAGE and Coomassie staining. The remaining cross-linked and non-crosslinked samples were separately pooled and precipitated using cold (−20 °C) acetone with overnight incubation. Dried protein pellets were resuspended in 12.5 μL of resuspension buffer [50 mM ammonium bicarbonate and 8 M urea (pH 8.0)]. ProteaseMAX (Promega, V5111) was added to a final concentration of 0.02%, and the solutions were mixed on an orbital shaker operating at 600 rpm for 5 min. After resuspension, 87.5 μL of digestion buffer [50 mM ammonium bicarbonate (pH 8.0)] was added. Protein solutions were digested using trypsin (Promega, V5280) at a ratio of 1:150 (w/w) (trypsin:protein) for 4 hours at 37°C, then digested overnight at room temperature using chymotrypsin (Promega, V1061) at a ratio of 1:150 (w/w) (chymotrypsin:protein). Peptides were acidified to a final concentration of 1% TFA and desalted using C18 ZipTip (Millipore Sigma, ZTC185096). Desalted peptides were dried using a vacuum centrifuge (SpeedVac, ThermoFisher). Dried peptides were resuspended in 10 μL of 0.1% TFA in water. Samples were frozen and stored at -20°C until LC-MS analysis.

#### Liquid chromatography and mass spectrometry

A total of 250 ng of sample was injected (triplicate injections for cross-linked samples and duplicate injections for control samples) onto an UltiMate 3000 UHPLC system (Dionex, ThermoFisher Scientific). Peptides were trapped using an Acclaim PepMap C18 trapping column (ThermoFisher Scientific) using a load pump operating at 10 μL/min. Peptides were separated on a 200-cm μPAC C18 column (PharmaFluidics/ThermoFisher) using a linear gradient (1% solvent B for 4 min, 1 to 30% solvent B from 4 to 70 min, 30 to 55% solvent B from 70 to 90 min, 55 to 97% solvent B from 90 to 112 min, and isocratic at 97% solvent B from 112 to 120 min) at a flow rate of 800 nL/min. Gradient solvent A contained 0.1% formic acid, and solvent B contained 80% acetonitrile and 0.1% formic acid. LC eluate was interfaced to an Orbitrap Fusion Lumos Tribrid mass spectrometer (ThermoFisher Scientific) with a Nanospray Flex ion source (ThermoFisher Scientific). The source voltage was set to 2.5 kV, and the S-Lens RF level was set to 30%. Cross-links were identified using a previously described MS2-MS3 method^70^ with slight modifications. Full scans were recorded from mass/charge ratio (m/z) 150 to 1500 at a resolution of 60,000 in the Orbitrap mass analyzer. The automatic gain control (AGC) target value was set to 4E5, and the maximum injection time was set to 50 ms in the Orbitrap. MS2 scans were recorded at a resolution of 30,000 in the Orbitrap mass analyzer. Only precursors with charge states between 4 and 8 were selected for MS2 scans. The AGC target was set to 5E4, a maximum injection time of 150 ms, and an isolation width of 1.6 m/z. Collision-induced dissociation fragmentation energy was set to 25%. The two most abundant reporter doublets from the MS2 scans with a charge state of 2 to 6, a 31.9721-Da mass difference, and a mass tolerance of ±10 parts per million (ppm) were selected for MS3. The MS3 scans were performed in the ion trap in rapid mode using higher-energy C-trap dissociation (HCD) fragmentation of 35% collision energy. The AGC target was set to 2E4, the maximum injection time was set to 200 ms, and the isolation width was set to 2.0 m/z.

#### Data analysis

To identify cross-linked peptides, Thermo *.raw files were imported into Proteome Discoverer 3.0 (ThermoFisher Scientific) and analyzed via XlinkX algorithm^71^ using the MS2_MS3 workflow with the following parameters: MS1 mass tolerance, 10 ppm; MS2 mass tolerance, 20 ppm; MS3 mass tolerance, 0.5 Da; digestion: trypsin/chymotrypsin with four missed cleavages allowed; minimum peptide length: 4; variable modification: oxidation (M), DSSO (K, S, T, and Y), and hydrolyzed DSSO (K, S, T, and Y). The XlinkX/PD Validator node was used for cross-linked peptide validation with a 1% false discovery rate. Identified cross-links were further validated and quantified using Skyline (version 19.1)^72, 73^ using a previously described protocol.^74^ Cross-link spectral matches found in Proteome Discoverer were exported and converted to sequence spectrum list format using Excel (Microsoft). Cross-link peak areas were assessed using the MS1 full-scan filtering protocol for peaks within 8 min of the cross-link spectral match identification. Peak areas were assigned to the specified cross-linked peptide identification if the mass error was within 10 ppm of the theoretical mass, the isotope dot product was greater than 0.90, and the peak was not found in the non–cross-linked negative control samples. The isotope dot product compares the distribution of the measured MS1 signals against the theoretical isotope abundance distribution calculated based on the peptide sequence. Its value ranges between 0 and 1, where 1 indicates a perfect match.^75^ Pairwise comparisons were made using the “MSstats” package implemented in Skyline to calculate relative fold changes and significance. Significant change thresholds were defined as a log2 fold change less than −1 or greater than 1 and −log_10_ P value greater than 1.3 (P value less than 0.05). Visualization of proteins and cross-links was generated using xiNET.^76^ Volcano plots were reformatted using Skyline output data within R with the tidyverse and ggplot2 packages.^77^

The data have been deposited to the ProteomeXchange Consortium via the PRIDE^69^ partner repository with the dataset identifier PXD056360.

### Structure Prediction

PR models and combination PR:SRC3:p300 ternary models were generated using five independent runs with different seeds using the AlphaFold3.0 server.^45^ The sequences provided were from the Uniprot P40601 isoforms for PR, Q9Y6Q9 for SRC3, and Q09472 for p300. The 32bp canonical PRE was modeled after the PRE used for crystallization experiments with the PR DBD^1^ (RCSB:2C7A). Top models from each independent run were opened and aligned in ChimeraX (version 1.8^78–80^). HDX-MS outputs were taken from HDXWorkbench^67^ and overlayed onto the generated models. Top models were selected based on average HDXer^46^ root mean square error (RMSE) values. HDXer was run using an automated server-based system to generate topology and gromacs files using the AMBER3.0 force fields. Topologies were then submitted to the server-based pipeline and RMSE values were generated from the differences between predicted and experimental deuterium exchange values.

### Progesterone receptor response assays

PR protein expression and response assays were co-transfected in 293T cells (ATCC, CRL-3216) at a 2:1 (protein:response plasmid) ratio using X-TremeGeneHP (Roche, XTGHP-RO) at a 4:1 (X-TremeGene:Plasmid Mix) ratio. Cells were incubated overnight, then re-seeded in 384-well plates in quadruplicate per transfected sample. Transfected cells were incubated for 4 hours at 37°C in a humidified 5% CO_2_ incubator. After 4 h incubation, either vehicle (EtOH), R5020 (Revvity, NLP004005MG), or RU486 (Sigma, M8046) were added 2:1 to the transfected cells for a final concentration of 50nM. Cells were incubated overnight with compound, then treated with Dual-Glo® reagents (Promega), following a standard protocol.^81^ Plates were read on a Bio-Tek Neo II plate reader using a gain of 200 and integration time of 100ms.

### Data Availability

Mass spectrometry data are available through the ProteomeXchange Consortium via the PRIDE partner repository under identifiers: PXD056400 and PXD056360. Interactive consolidated HDX-MS data are available through the hdxms.app web server using the following link: XXX All other datasets generated and/ or analyzed during the current study are available from the corresponding author upon reasonable request.

## Acknowledgements

This work was supported by NIH-NCI R01 (CA263574) to MPIs, PRG, RK, and DPE. The authors acknowledge the expert assistance of the Recombinant Protein Production and Characterization Core (RPPCC) for expression of recombinant proteins in the baculovirus insect cell system and the Mass Spectrometry Proteomics (MSPC) Core for amino acid sequencing and phosphorylation analysis of recombinant proteins. The RPPCC and MSPC at BCM are supported by the NCI Cancer Center Support Grant (P30 CA125123) of the Dan L Duncan Comprehensive Cancer Center and the MSPC is additionally supported by a CPRIT (Cancer Prevention and Research Institute of Texas) Core Facility Support Award (RP210227). Additional support is from NIH S10 Shared Instrument Grant (1S10OD030276) to JCF for the SEC-MALS instruments.

## Extended Data

**Extended Data Fig. 1.**
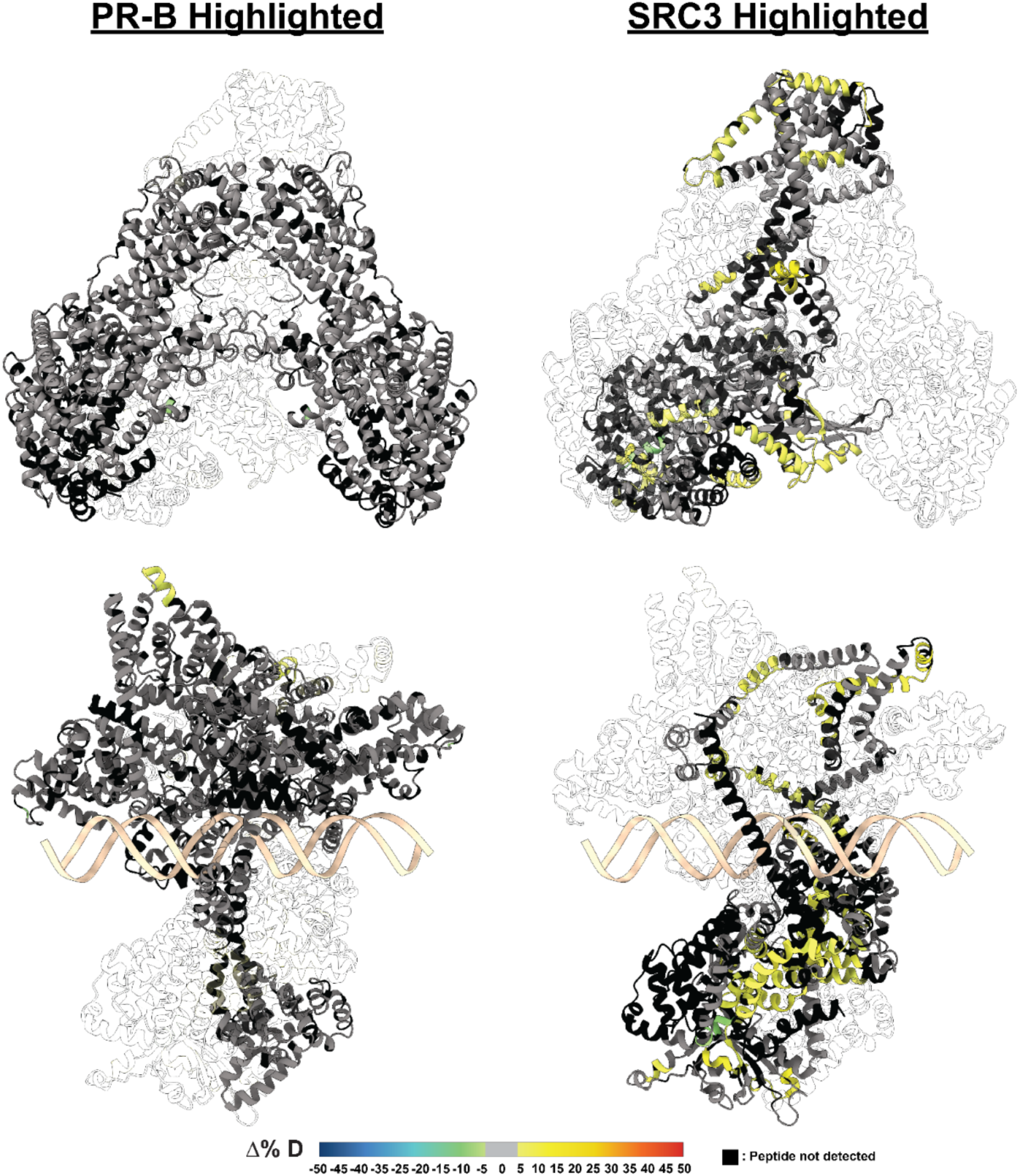
HDX overlays on best HDXer models of PR-B:SRC3 complexes. HDX-MS overlays on best selected model of PR-B:SRC3 complex in both non-DNA-bound (top) and DNA-bound (bottom) states. Warmer colors (yellow, orange, red), indicate increased deuterium exchange, and cooler colors (green, blue) indicate decreased deuterium exchange. Black indicates the peptide was not found by HDX and gray indicates exchange within ±5%. Models selected by HDXer RMSE values, and visualizations made using ChimeraX 1.8.

**Extended Data Fig. 2.**
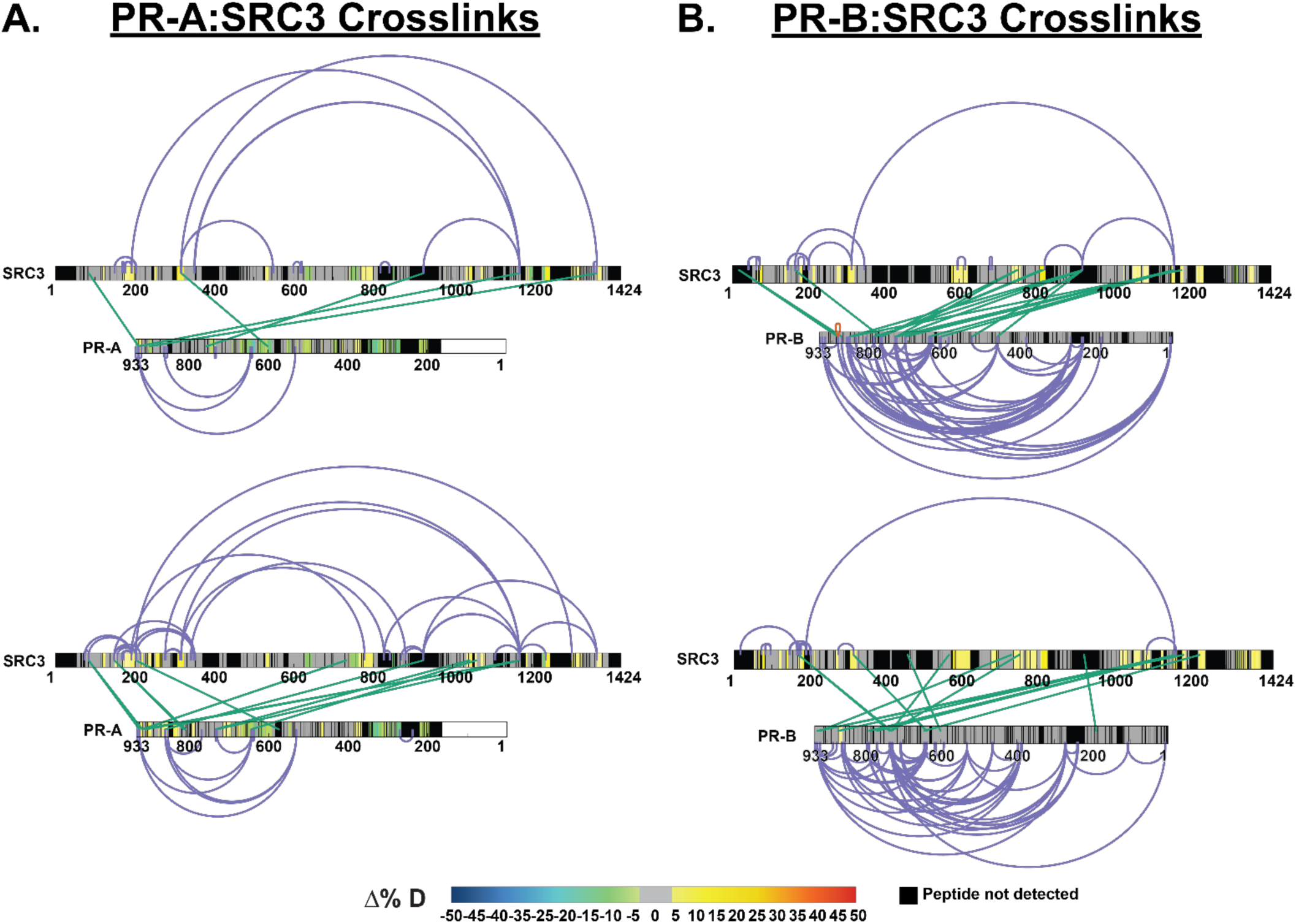
PR and SRC3 deuterium exchange differences align with crosslinking results. **A.** XiView images of differential XL-MS data for PR-A vs. PR-A:SRC3 experiments with crosslinks enriched in apo samples (top) and pre-bound samples (bottom) represented. HDX overlays are indicative of their comparable differential HDX-MS experiments. Top: SRC3 exchange - SRC3 vs. PR-A:SRC3 (non-DNA bound), PR exchange – PR-A vs. PR-A:SRC3.Bottom: SRC3 exchange – SRC3 vs. PR-A:SRC3:PRE, PR exchange – PR-A:SRC3 vs. PR-A:SRC3:PRE. **B.** XiView images of differential XL-MS data for PR-B vs. PR-B:SRC3 experiments with the same HDX overlays as shown in **A.** Intraprotein crosslinks are represented in purple, interprotein crosslinks are represented as green. Differential HDX-MS scale shows the % change in deuterium incorporation, where cooler colors represent reduced deuterium exchange, while warmer colors (yellow, orange, red) represent increased deuterium exchange. Black represents peptides not detected in HDX-MS experiments, and PR isoforms have numbering normalized to the B isoform.

**Extended Data Fig. 3.**
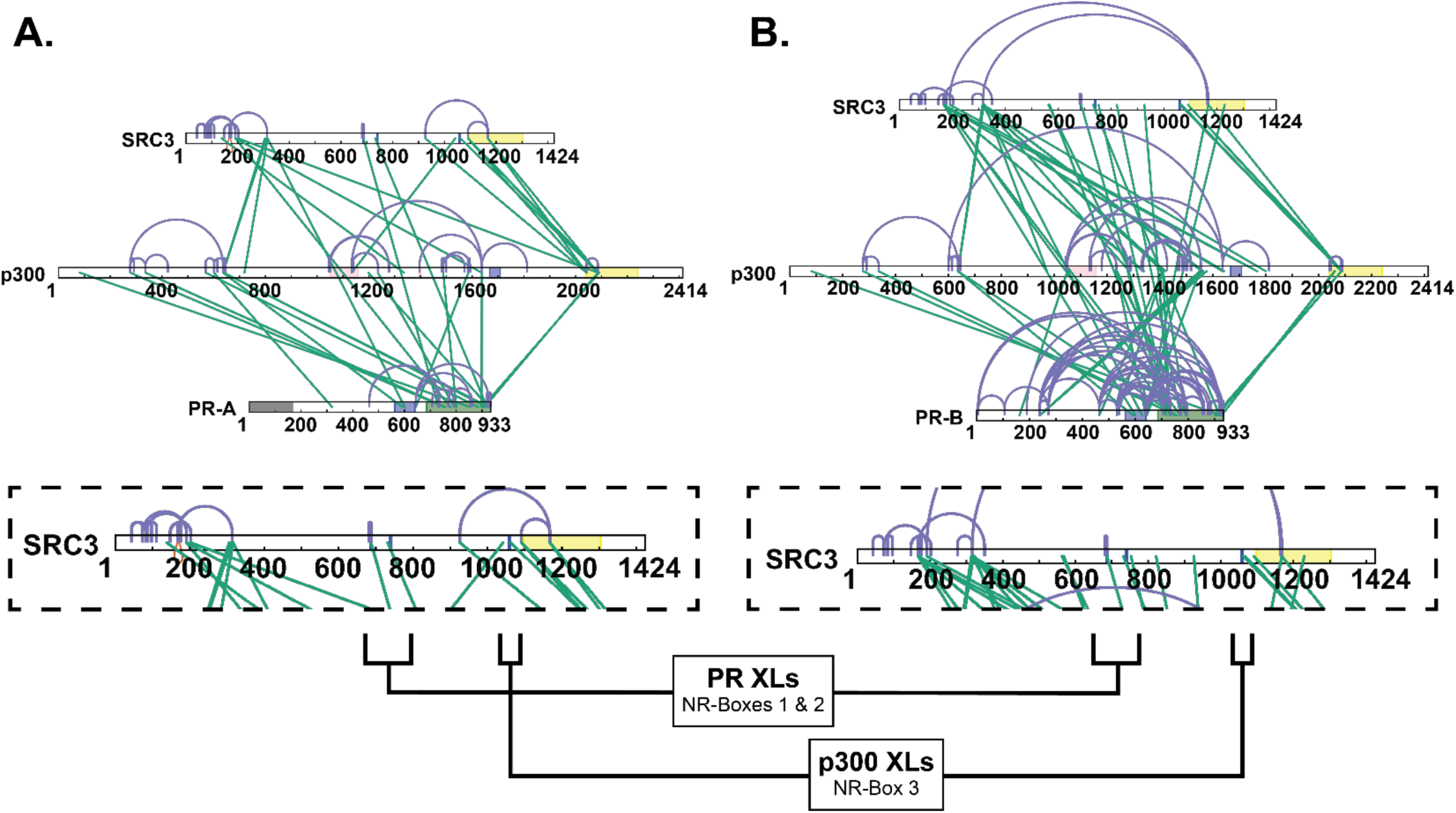
Crosslinking shows NR-Box 3 is exclusively used by p300. **A.** Top: All validated crosslinks from differential PR-A:SRC3:p300±PRE experiments. Bottom: Zoomed in view of SRC3 to show NR-Box crosslinks specific to the PR-A-containing complex. **B.** Top: All validated crosslinks from differential PR-B:SRC3:p300±PRE experiments. Bottom: Zoomed in view of SRC3 to show NR-Box crosslinks specific to the PR-B-containing complex. Defined domains are as follows: PR - DBD (purple) and LBD (green); SRC3 – NR-boxes (purple) and histone acetyltransferase domain (yellow); p300 - bromodomain (pink), zinc finger domain (green), and NCOA2-interaction domain (yellow).

**Extended Data Fig. 4.**
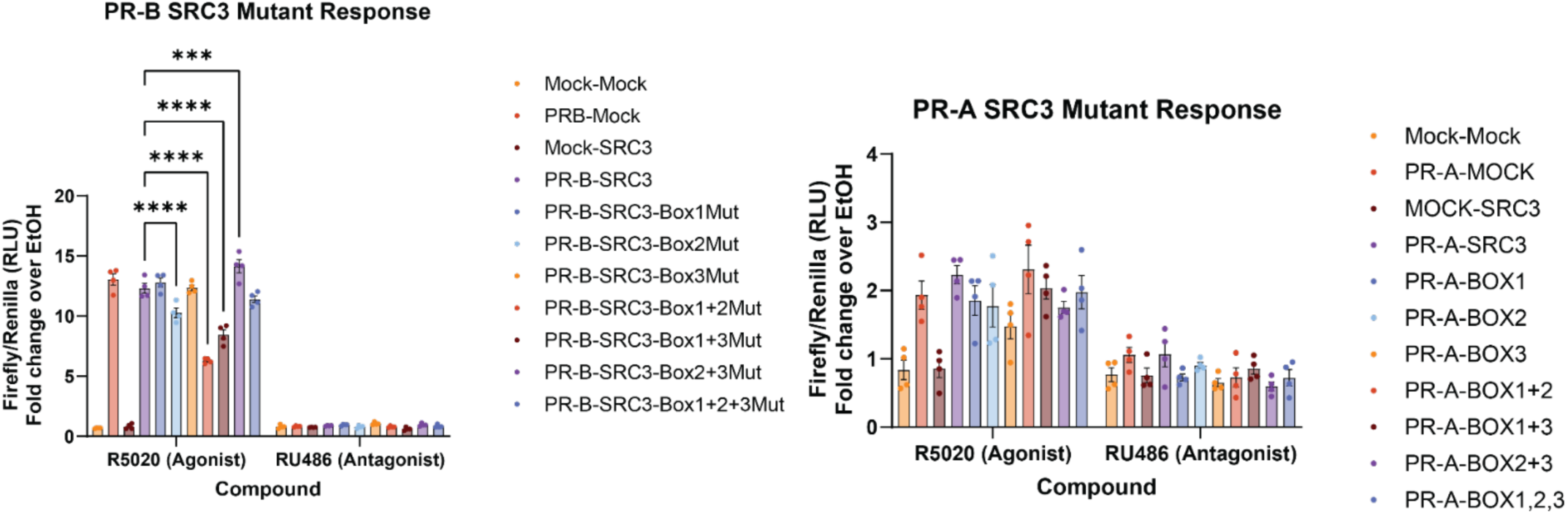
Mutation of NR-box 2 or multiple NR-boxes to LXXAA in SRC3 reduces PR activity. Bar graph showing the ratio of Firefly to Renilla luciferase normalized over the vehicle (ethanol) signal for both R5020 PR agonist and RU486 PR antagonist. Mock is indicative of base vector in the place of PR or SRC3 plasmid. ‘Mut’ in the legend indicates an LXXLL NR-Box mutated to LXXAA, validated by whole plasmid next-generation sequencing (Azenta). Similarly, ‘BOX’ denoted in the PR-A experiments indicates the mutated NR-Box(es). Points indicative of 4 replicates with SEM error bars. Two-way ANOVA was performed with Tukey correction for multiple comparisons, and asterisks indicate p&0.001 (***) and p&0.0001 (****).

**Extended Data Fig. 5.**
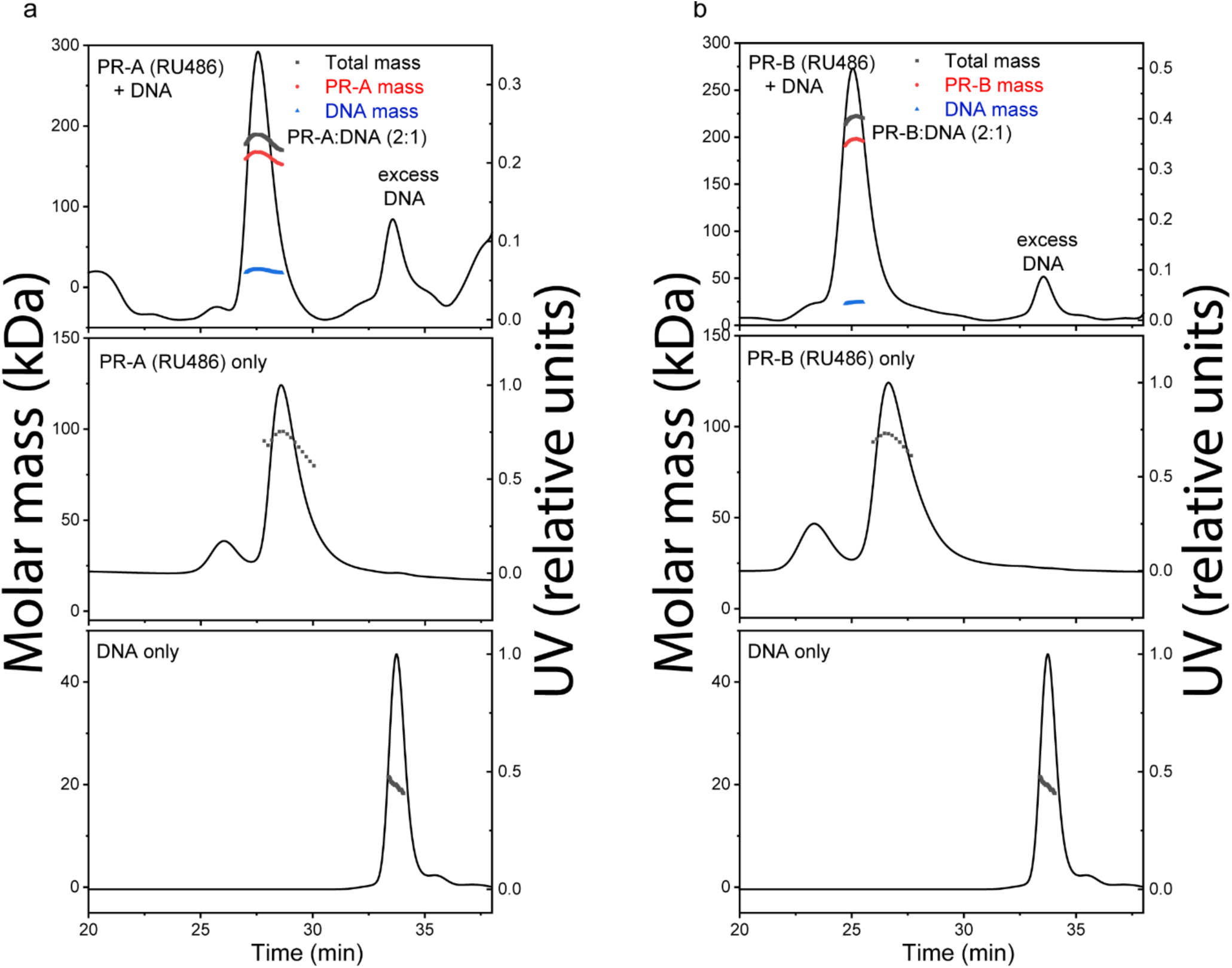
SEC-MALS of purified PR-A and PR-B bound to the progestin antagonist RU486 with and without DNA. DNA induces assembly of PR dimers A-B. SEC-MALS chromatograms of antagonist (RU486)-bound PR-A (A) and PR-B (B) with and without DNA. The molar mass of DNA and PR-A alone matches the monomeric molar mass (black line/dots across the peaks). DNA induces assembly of both PR-A and PR-B into a complex with 2:1 (protein:DNA) stoichiometry. The presence of DNA in the complexes were confirmed by deconvolution of the protein and DNA fractions in the peak (red and blue lines, respectively).

