## Supplemental for "Structural proteomics defines a sequential priming mechanism for the progesterone receptor"

Running title: Structural proteomics reveals a unique activation model for PR-A and PR-B

Acknowledgements: This work was supported by NIH-NCI R01 (CA263574) to MPIs, PRG, RK, and DPE. The authors acknowledge the expert assistance of the Recombinant Protein Production and Characterization Core (RPPCC) for expression of recombinant proteins in the baculovirus insect cell system and the Mass Spectrometry Proteomcs (MSPC) Core for amino acid sequencing and phosphorylation analysis of recombinant proteins. The RPPCC and MSPC at BCM are supported by the NCI Cancer Center Support Grant (P30 CA125123) of the Dan L Duncan Comprehensive Cancer Center and the MSPC is additionally supported by a CPRIT (Cancer Prevention and Research Institute of Texas) Core Facility Support Award (RP210227). Additional support is from NIH S10 Shared Instrument Grant (1S10OD030276) to JCF for the SEC-MALS instruments.

Keywords:
Progesterone receptor; hydrogen-deuterium exchange; crosslinking; mass spectrometry; protein-protein interactions; nuclear receptors, transcriptional co-regulatory proteins

**Supplemental Figures**

1. Affinity/SEC two-step purification of SII-tagged PR
2. Mass Spectrometry sequencing of purified major PR-A and PR-B bands
3. Mass Spectrometry sequencing of minor smaller molecular sized band product with PR-B
4. Affinity/SEC two-step purification for SRC3 tagged SII
5. Affinity/SEC two-step purification for p300 tagged SII
6. Mass Spectrometry of purified SRC3 and p300
7. SEC MALS of PR-A/SRC3/p300/DNA Complex
8. Differential scanning fluorimetry (DSF) analysis of PR-A with and without DNA
9. Intrinsic deuterium exchange for PR-A and PR-B
10. Workflow for PR:SRC3 model generation and selection
11. PR-B vs. PR-B:PRE consolidated HDX-MS data

**Supplemental Tables**

1. SEC-MALS Experimental vs Theoretical MWs of individual purified proteins and various mixtures in the presence and absence of DNA
2. Transition melting temperatures ($T_M$) of each purified protein as determined by DSF

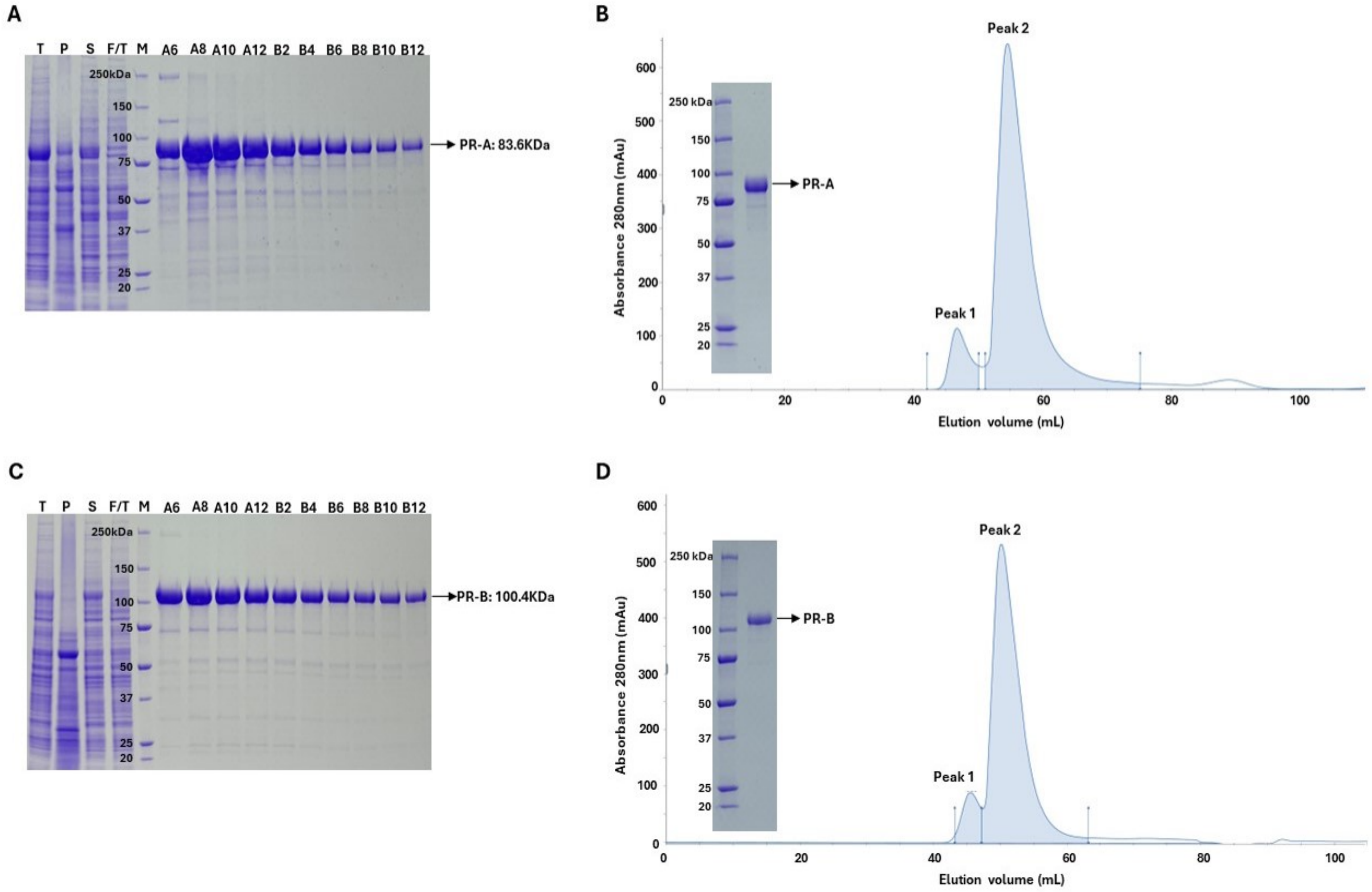


**Fig S1. Affinity/SEC two-step purification of SII-tagged PR.** SII-tagged PR-A and PR-B(bound to progestin agonist R5020) were expressed in Sf9 cells and cell lysates were prepared and purification performed by two -step affinity Strep Trap XT column and size exclusion chromatography (SEC) S200 as described in Methods. A) SDS-PAGE analysis of PR-A affinity purification fractions T, total cell lysate, P pellet after centrifugation lysate, S supernatant after centrifugation, F/T flow through after binding to Strep TrapXT, M, protein molecular weight markers. Lanes A6-B12 (5uL) biotin eluted fractions with major enriched protein bands at expected size of ~83 kDa for PR-A. B) SEC fractionation of pooled fractions A6-B12 off the Strep Trap XT column. OD tracing at 280nM showsmajor peak 2 of monomeric size for PR-A and peak 1 containing high molecular weight aggregation. Peak 2 fractions were pooled, concentrated and analyzed SDS-PAGE gel showing a single band of >98% purity of the expected size of PR-A. C) A) SDS-PAGE analysis of PR-B affinity purification fractions and D) SEC second step purification by the same set-up and as PR-A except major band and purified product are expected size for monomeric PR-B (~100kDa).

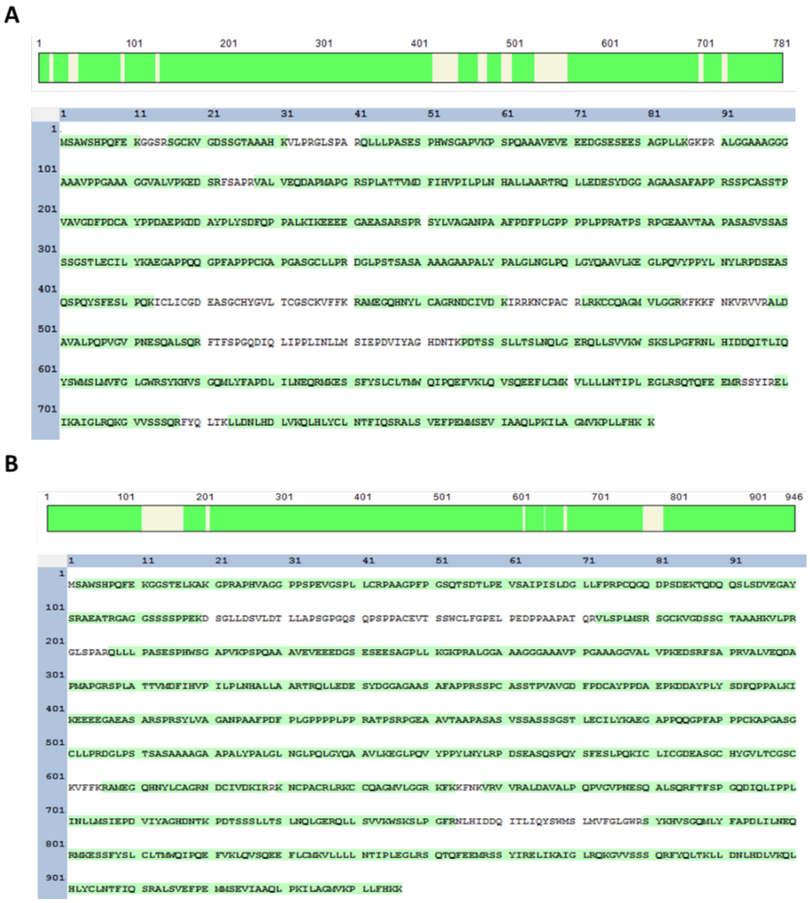

**Fig S2. Mass Spectrometry sequencing of purified major PR-A and PR-B bands.** In-gel digestion of purified major bands for A) PR-A and B) PR-B (bound to progestin agonist R5020) was performed, followed by LC-MS/MS analysis. The peptide sequences highlighted in green were identified by MS, covering nearly the entire tryptic region

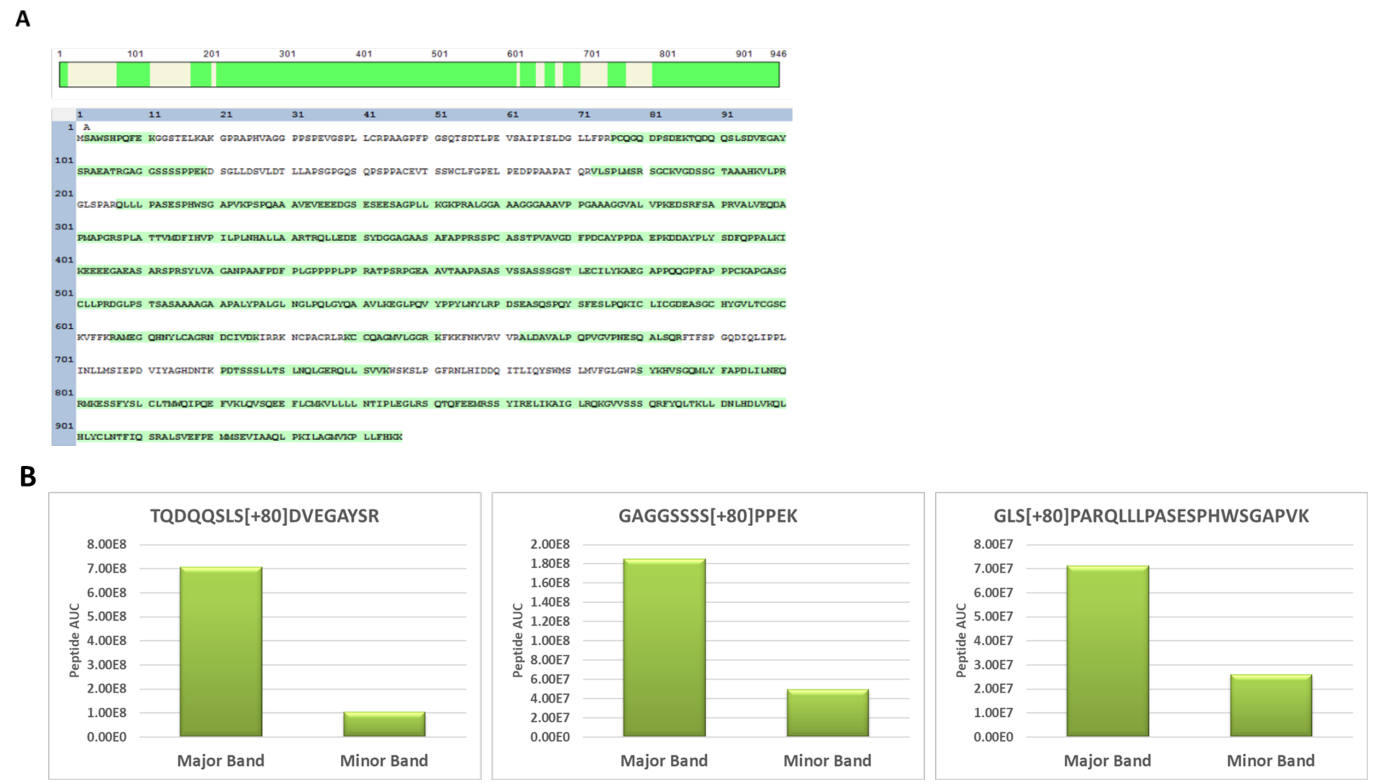


**Fig S3. Mass Spectrometry sequencing of minor smaller molecular sized band product with PR-B.** A) In-gel digestion of smaller sized PR-B band (bound progestin agonist R5020) was performed, followed by LC-MS/MS analysis. The peptide sequences highlighted in green were identified by MS, suggesting intact full-length PR-B protein. B) The peptide area under curve (AUC) for selected phosphorylated PR-B peptides from major and minor band suggests variable phosphorylation level. The peptide AUC was normalized to the total protein amount in that band.

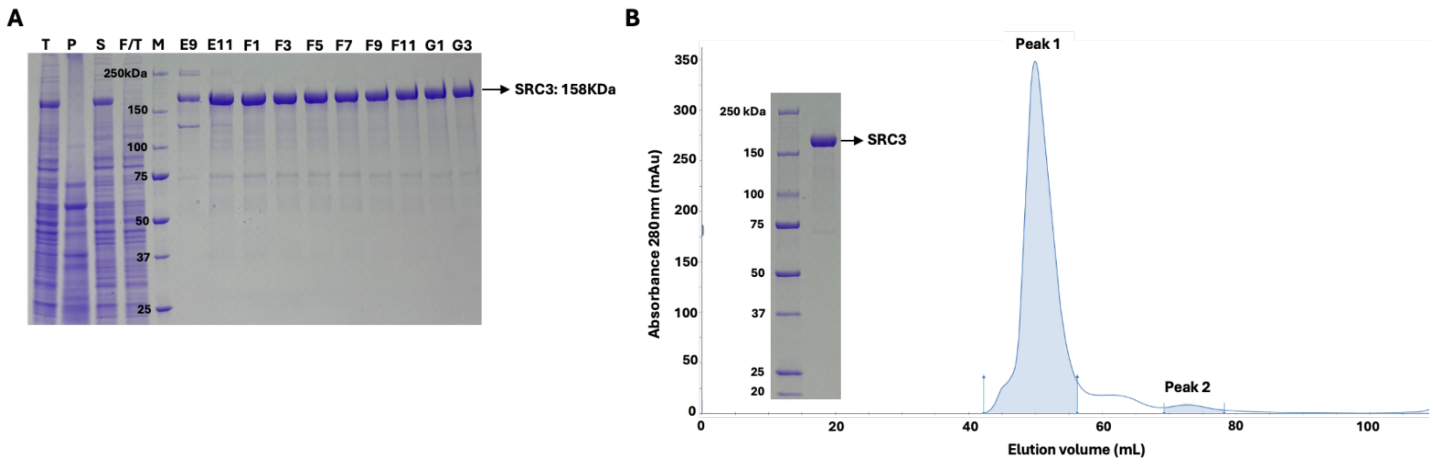


**Fig S4. Affinity/SEC two-step purification for SRC3 tagged SII.** SII-tagged SRC3 was expressed in Sf9 insect cells and purification performed by two -step affinity by StrepTrap XT and size exclusion chromatography (SEC) S200 as described in Methods. A) SDS-PAGE analysis of affinity purification fractions T, total cell lysate, P pellet after centrifugation, S supernatant after centrifugation, F/T flow through after binding to Strep TrapXT M, protein molecular weight markers. Lanes E9-G3 are SDS-PAGE analysis (5uL) of biotin eluted fractions with a major enriched band of expected size of ~158kDa for SRC3. B) SEC (S200) fractionation of pooled fractions E11-G3 off the StrepTrap XT column. OD tracing at 280nM shows a major peak (peak 1) of the native size expected for monomeric SRC3 and a minor peak (peak 2) of lower molecular weight material. Peak 1 fractions were pooled, concentrated and analyzed by SDS-PAGE(left panel) showing a single band of >98% purity of the expected size of monomeric SRC3

.

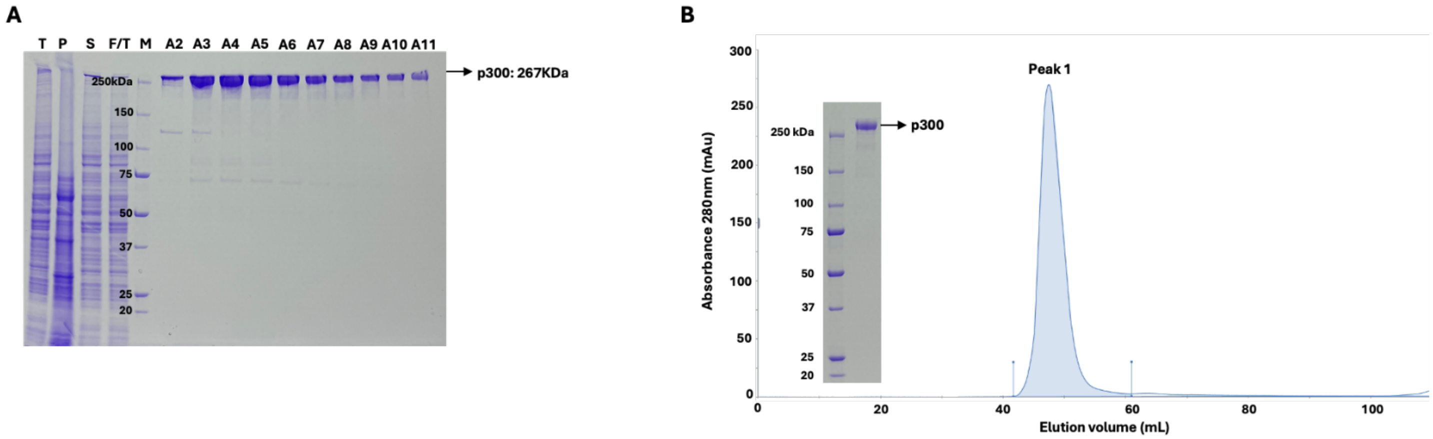


**FigS5 Affinity/SEC two-step purification for p300 tagged SII.** SII-tagged p300 was expressed in Sf9 insect cells and purification performed by two -step affinity by StrepTrap XT and size exclusion chromatography (SEC) S200 as described in Methods. *A)* SDS-PAGE analysis of affinity purification fractions T, total cell lysate, P pellet after centrifugation, S supernatant after centrifugation, F/T flow through after binding to Strep TrapXT M, protein molecular weight markers. Lanes A2-A11 are SDS-PAGE ( 5uL) of biotin eluted fractions with a major enriched band of the expected size for monomeric p300 (~267kDa). *B)* SEC (S200) fractionation of pooled fractions A3-A8 from Strep TrapXT shows a single major peak (OD tracing at 280nM) of expected size for native monomeric p300. The pooled SEC fractions from peak 1 on SDS-PAGE (left panel) analysis shows a single major band of the expected sized for monomeric p300.

A

B

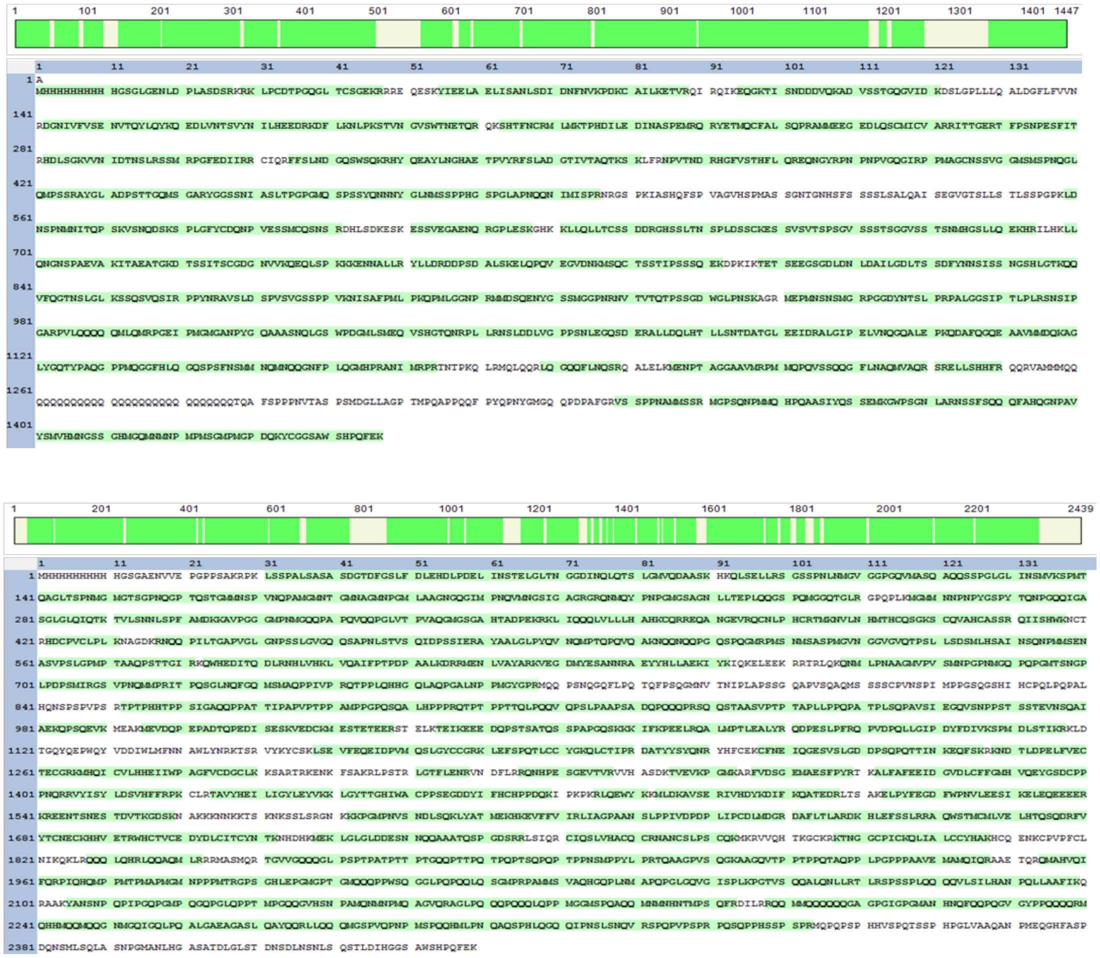

**Fig S6. Mass Spectrometry of purified SRC3 and p300.** In-gel digestion of purified A) SRC3 and B) p300 bands was performed, followed by LC-MS/MS analysis. The peptide sequences highlighted in green were identified by MS, covering nearly the entire tryptic region.

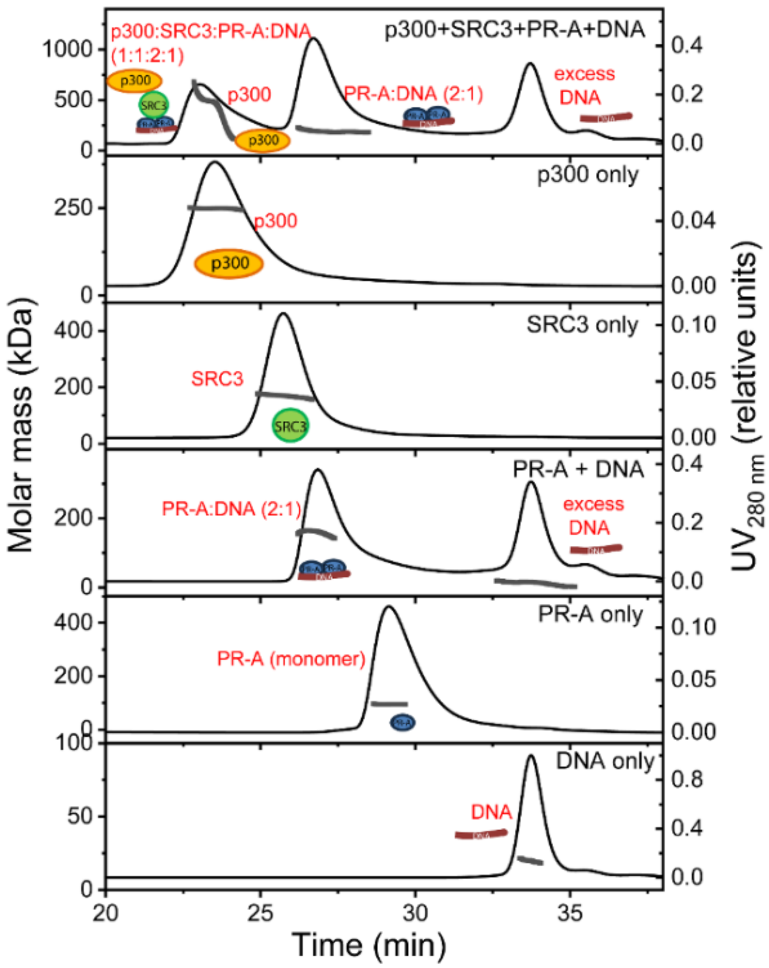


**Fig S7 SEC-MALS chromatograms with molar mass distribution for various proteins (PR-A, SRC3, p300) and DNA alone or in complexes.** Shown are double y-plots (left y-axis, molar mass; right y-axis, UV absorbance at 280 nm) vs time (x-axis). Molar mass distribution is displayed as gray dots across the peaks. PR-A (with agonist R5020) assembles only to a dimer in the presence of DNA. A multi-complex of p300:SRC3:PR-A:DNA at a 1:1:2:1 ratio is observed (top chromatogram). The presence of DNA in the complex is confirmed by deconvolution of the protein and DNA fractions in the peak. The full complex showed a heterogeneous distribution of molar mass due to overlap with the p300 monomeric peak.

**Table S1. SEC-MALS Experimental vs Theoretical MWs of individual purified proteins and various mixtures in the presence and absence of DNA.**

Table 1. SEC-MALS derived molecular weights for proteins, DNA and the complexes formed

| | Theoretical MW | Actual MW[a] | |
|---|---|---|---|
| Protein/DNA | | by UV | by dRI |
| DNA | 19.6 | 18.2 ± 0.9 | 20.1 ± 1.0 |
| PR-A(R5020) | 83.6 | 94.9 ± 4.8 | 91.2 ± 4.6 |
| PR-A(RU486) | 83.6 | 98.8 ± 4.7 | 95.3 ± 4.8 |
| PR-B(R5020) | 100.4 | 116.4± 5.8 | 101.6± 5.1 |
| PR-B(RU486) | 100.4 | 96.3 ± 4.8 | 97.7 ± 4.10 |
| p300 | 267 | 249.2 ± 12.5 | 200.5 ± 9.2 |
| SRC3 | 158 | 170.3 ± 8.5 | 159.7 ± 8 |

| Complexes | Theoretical MW | Actual MW |
|---|---|---|
| PR-A(R5020):DNA (2:1) | 186.8 | 161.6 ± 8.1 |
| Protein component | 167.2 | 144.8 ± 7.2 |
| DNA component | 19.6 | 16.8 ± 0.8 |
| PR-A(RU486):DNA (2:1) | 186.8 | 189.3 ± 9.5 |
| Protein component | 167.2 | 167.5 ± 8.4 |
| DNA component | 19.6 | 21.8 ± 1.1 |
| PR-B(R5020):DNA (2:1) | 220.4 | 224.0 ± 11.2 |
| Protein component | 200.8 | 201.7± 10.1 |
| DNA component | 19.6 | 22.3 ± 1.1 |
| PR-B(RU486):DNA (2:1) | 220.4 | 221.8 ± 11.1 |
| Protein component | 200.8 | 197.7± 9.9 |
| DNA component | 19.6 | 24.1 ± 1.2 |
| p300:SRC3:PR-A(R5020):DNA (1:1:2:1) | 611.8 | 536.5 ± 26.8[b] |
| Protein component | 592.2 | 512.7 ± 25.6 |
| DNA component | 19.6 | 23.7 ± 1.2 |
| PR-A(R5020):DNA (2:1)[c] | 186.8 | 190.7 ± 9.5 |
| Protein component | 167.2 | 161.4 ± 8.1 |
| DNA component | 19.6 | 29.3 ± 1.5 |

[a] errors are 5% of calculated MW; dn/dc based off BSA values in similar buffer condition

[b] heterogeneous MW distribution, overlap with p300

[c] in same buffer conditions as above but with 0M Urea

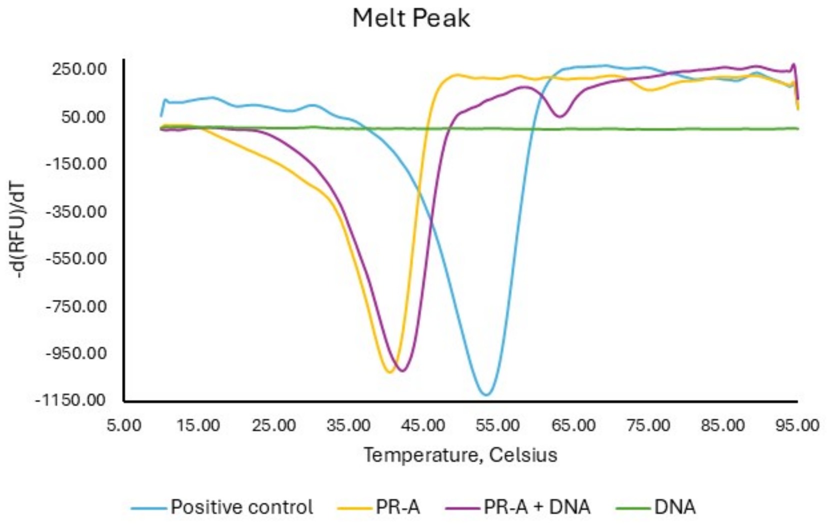


**Fig S8. Differential scanning fluorimetry (DSF) analysis of PR-A with and without DNA.** DSF analysis was performed with 4 µM PR-A and 25× SYPRO Orange in 20 mM HEPES, pH 7.5, 200mM NaCl, 5% Glycerol, 1M Urea, 1mM TCEP with (purple) or without 3uM PRE DNA (yellow), compared with the same concentration of TLX-WT as a positive control (blue). Fluorescent intensities and the derivative of the fluorescent intensities (dF/dT) were plotted against temperature (Tm).

**Table S2. Transition melting temperatures ($T_M$) of each purified protein as determined by DSF.** Samples were analyzed as three technical replicates and the average Tm was calculated with standard error of the mean.

| Protein | $T_m$ (°C) |
|---|---|
| PR-A | 40.7±0.3 |
| PR-A + DNA | 42.2±0.3 |
| PR-B | 43.5±0 |
| PR-B + DNA | 45.0±0 |
| p300 | 49.1±0 |
| SRC3 | 46.5±0.1 |

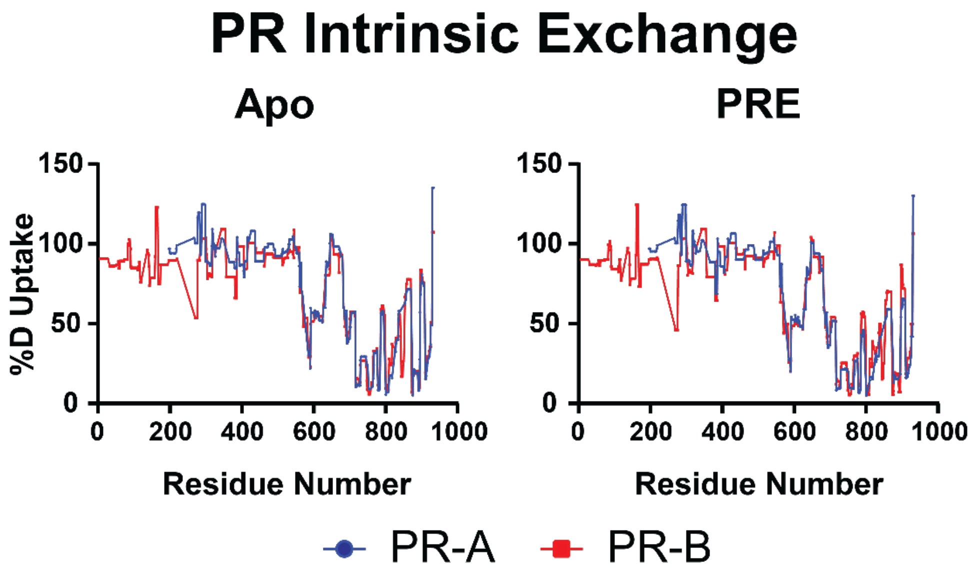


**Figure S9**. **Intrinsic deuterium exchange for PR-A and PR-B**. Woods plots showing the percent deuterium exchange (%D Uptake) for both PR-A (blue) and PR-B (red) across each residue across the protein in both non-DNA-bound (Apo) and DNA-bound (PRE) states. Graphs made using GraphPad Prism 10.

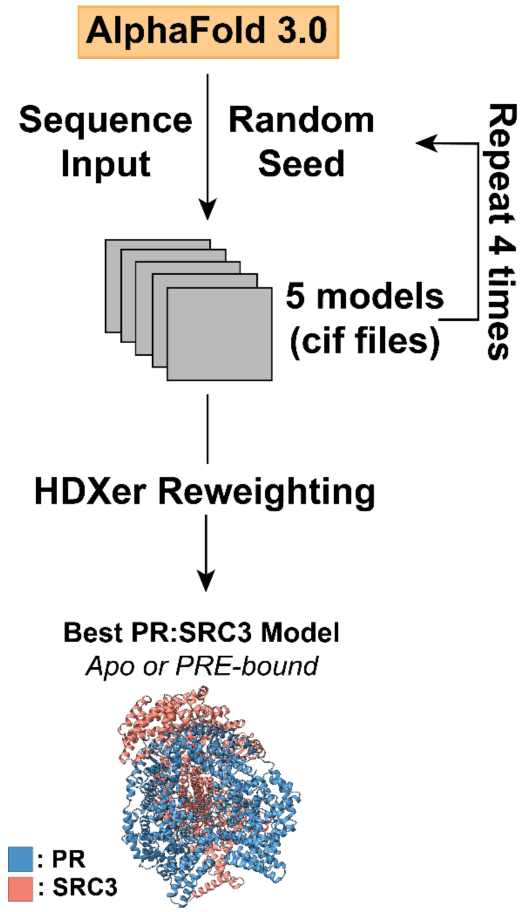


**Figure S10 Workflow for PR:SRC3 model generation and selection**. Workflow detailing the model generation for ternary complexes. AlphaFold3.0 webserver was used to generate PR:SRC3 models in quintuplicate. Models were ensemble reweighted using HDXer (see methods) to determine best representation based on HDX data.

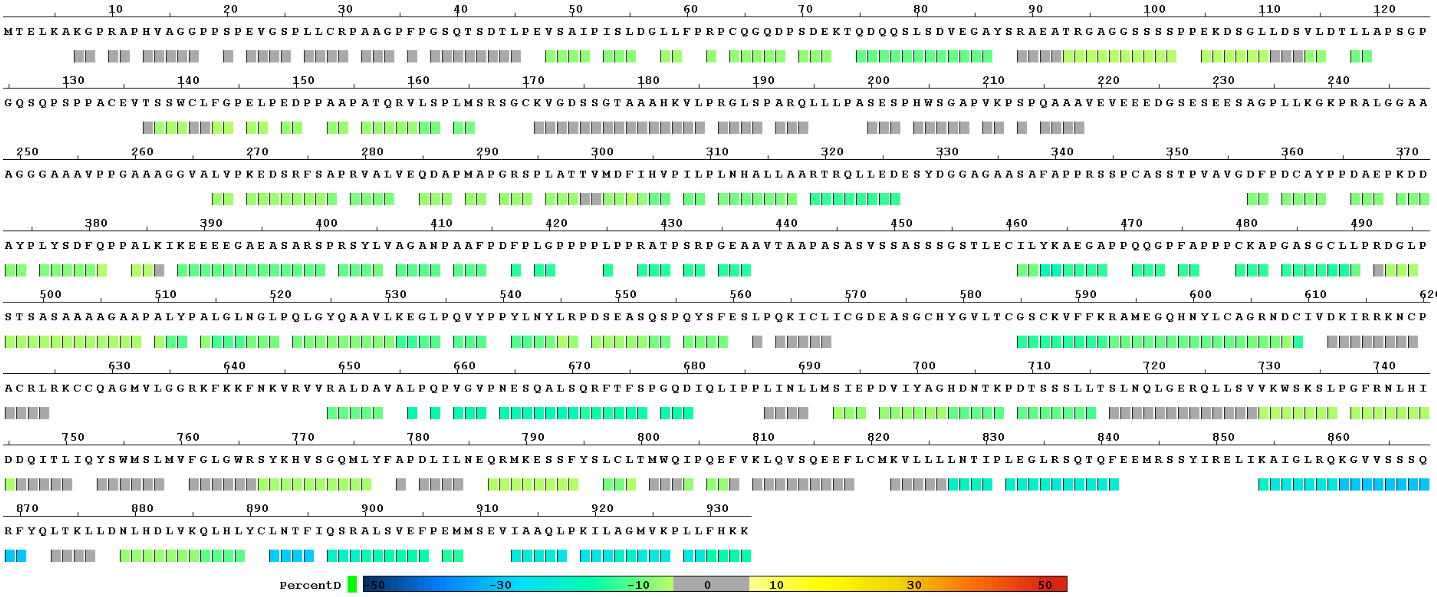


**Figure S11 PR-B vs. PR-B:PRE consolidated HDX-MS data.** *Consolidated data output from HDXWorkbench for PR-B ± DNA HDX-MS experiments when RU-486 bound. This illustrates the concerted decrease in deuterium exchange when DNA bound, compared to non-DNA-bound protein. Each point represents an average across each peptide detected, with the differential HDX data following the color scheme shown at the bottom.*
